# Estimating Brain Similarity Networks with Diffusion MRI

**DOI:** 10.1101/2025.03.29.646134

**Authors:** Amir Sadikov, Hannah L. Choi, Lanya T. Cai, Pratik Mukherjee

**Affiliations:** Radiology and Biomedical Imaging, University of California, San Francisco; Graduate Group in Bioengineering, University of California, San Francisco

**Keywords:** Diffusion MRI, Structural Covariance, Morphometric Similarity, Brain Networks, Connectome, Gray Matter Microstructure

## Abstract

Structural similarity has emerged as a promising tool in mapping the network organization of an individual, living human brain. Here, we propose diffusion similarity networks (DSNs), which employ rotationally invariant spherical harmonic features derived from diffusion magnetic resonance imaging (dMRI), to map gray matter structural organization. Compared to prior approaches, DSNs showed clearer laminar, cytoarchitectural, and micro-architectural organization; greater sensitivity to age, cognition, and sex; higher heritability in a large dataset of healthy young adults; and straightforward extension to non-cortical regions. We show DSNs are correlated with functional, structural, and gene expression connectomes and their gradients align with the sensory-fugal and sensorimotor-association axes of the cerebral cortex, including neuronal oscillatory dynamics, metabolism, immunity, and dopaminergic and glutaminergic receptor densities. DSNs can be easily integrated into conventional dMRI analysis, adding information complementary to structural white matter connectivity, and could prove useful in investigating a wide array of neurological and psychiatric conditions.

## Introduction

Recent advances in magnetic resonance imaging (MRI) analysis have enabled mapping of brain similarity networks for individual, living human brains [Sebenius et al., 2025]. Previously, structural covariance networks (SCNs), computed via subject-wise correlation of a metric, such as cortical thickness, for every pair of regions, could produce an aggregate similarity network, but lacked individual specificity [Alexander-Bloch et al., 2013; Lerch et al., 2006; Zielinski et al., 2010]. Morphometric similarity networks (MSNs), which measured similarity via pairwise correlation of standardized feature vectors, were able to produce individual brain networks, but suffered from two key limitations: they reduced the rich spatial information of structural MRI (sMRI) to a single summary statistic and forced features to have equivalent variance for every region [Seidlitz et al., 2018]. Morphometric inverse divergence (MIND) networks, which computed similarity between gray matter regions based on the symmetric Kullback-Leibler (KL) divergence of their multivariate distributions of structural features, were found to be more robust, more consistent with cortical cytoarchitecture, better correlated with tract tracing measures of axonal connectivity, more sensitive to age-related changes, and more heritable over edge weight and degree centrality statistics [Sebenius et al., 2023].

Despite the improvement of MIND over previous similarity measures, MIND suffers from two limitations inherent to its usage of sMRI. First, the structural features that MIND uses, namely cortical thickness, surface area, volume, mean curvature, and sulcal depth, can only be estimated in the cortex and so extension to subcortical regions is not possible. Second, while morphological features derived from sMRI can be informative, they lack the rich information that can be gleaned from other MRI contrasts, such as diffusion MRI (dMRI), which directly interrogates tissue microstructure by mapping the diffusion of water [Jones, PhD, 2012; Novikov et al., 2016]. For instance, high resolution ex-vivo maps of dMRI signal representations, such as diffusion tensor imaging (DTI) [Basser et al., 1994] and higher-order mean apparent propagator (MAP-MRI) [Özarslan et al., 2013], were able to reveal laminar substructures and were correlated with histological markers of cytoarchitecture [Avram et al., 2022; Wang et al., 2020]. In addition, in-vivo cortical dMRI could distinguish Mesulam’s hierarchy of laminar differentiation and von Economo and Koskinas cytoarchitectonic classes [Fukutomi et al., 2018; Sadikov et al., 2024]. The precision and sensitivity of dMRI to tissue microstructure make it a promising candidate for generating similarity networks.

Most work to date on brain similarity networks has focused solely on cortico-cortical similarity [Homan et al., 2019; Joo et al., 2024; Morgan et al., 2019; Qu et al., 2024; Tranfa et al., 2024; Wang et al., 2024; Xue et al., 2023; Zong et al., 2023]. A recent publication has extended MSNs to other brain structures using a wide range of metrics, including subcortical volume, T1w and T2w signal, and dMRI microstructural metrics from DTI and restriction spectrum imaging (RSI), to relate subcortical-cortical dissimilarity in preadolescents to cognitive function and psychiatric symptomology [Wu et al., 2023]. Apart from this article, there is limited literature exploring subcortical-cortical or subcortical-subcortical similarity noninvasively or using dMRI signal features to compute structural similarity between gray matter regions.

We introduce diffusion similarity networks (DSNs), which extend the MIND framework to dMRI. For DSNs, we use rotationally invariant spherical harmonic (RISH) [Kazhdan et al., 2003] descriptors instead of macrostructural features, and perform our computations voxel-wise as opposed to vertex-wise. These two changes allow us to extend structural similarity to any region of interest. RISH features measure the energies of the frequency components of a spherical function and represent the optimal linear decomposition of a spherical function into rotationally invariant components. RISH features are robust to noise, flexible to a wide array of spherical sampling schemes, and are widely used in dMRI analysis from fiber orientation estimation [Anderson, 2005; Descoteaux et al., 2007; Hess et al., 2006] and multi-site scanner harmonization [Cetin Karayumak et al., 2019; Mirzaalian et al., 2018] to biophysical model fitting [Coelho et al., 2022]. Importantly, RISH decomposition can be generally applied to any spherical signal and, unlike microstructural metrics, does not make any underlying assumptions about the dMRI signal.

In a large healthy young adult cohort, we compared DSNs to MSNs, MIND networks, structural connectivity (SC) computed via dMRI tractography [Hubbard and Parker, 2013; Mukherjee et al., 2008], and functional connectivity (FC) [Biswal et al., 1995; Lee et al., 2013] computed via resting-state functional MRI (rs-fMRI) across the following evaluation metrics: 1) reliability, measured via between-subject variability, intra-subject test-retest reproducibility, and intra-class correlation coefficients (ICC); 2) biological validity, measured via adherence to known principles of anatomical organization, such as cytoarchitecture [Constantin von Economo et al., 1925; Scholtens et al., 2018], laminar differentiation [Mesulam, 1998], and micro-architecture [Glasser and Van Essen, 2011]; and 3) sensitivity, measured via prediction of age, sex, cognition as well as via heritability of edge and node attributes in network analysis. We replicate our results in an independent ultra-high b-value dMRI dataset [Fan et al., 2016], demonstrating consistent biological fidelity. Additionally, we show that DSNs can be accurately computed from a subsampled single shell dMRI acquisition (b=1000 s/mm^2^; 30 directions) [Jones, 2004] and retain many of the same advantages as DSNs derived from multi-shell dMRI acquisitions. With DSNs, we also investigate subcortical-cortical and subcortical-subcortical homophily in the living human brain [Akarca et al., 2022; Alexander-Bloch et al., 2013; Goulas et al., 2019; Kerstjens et al., 2022; Oldham et al., 2022]. DSNs can be easily computed, applied to subcortical areas, add information that is complementary to structural connectivity computed using dMRI tractography, and could prove useful in investigating a wide variety of neurological and psychiatric conditions.

## Materials and Methods

### DSN Estimation

DSN estimation follows the same steps as MIND estimation with two caveats: we choose to use RISH features instead of morphometric features, and we compute our statistics voxel-wise as opposed to vertex-wise. The diffusion-weighted signal for each spherically sampled shell can be linearly decomposed onto the orthonormal spherical harmonics:

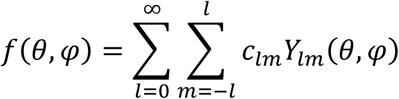

RISH features can then be measured by computing the energy of the frequency components of each shell as follows:

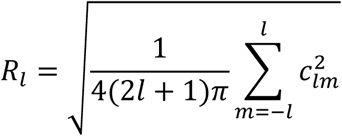

We estimated the spherical coefficients *c*_*lm*_ for each shell, up to the 4^th^ degree, via least-squares fitting with Laplacian regularization of 0.006 to prevent overestimation of higher frequency shells [Descoteaux et al., 2006]. Since diffusion weighting is inherently symmetric, we only estimate the coefficients for even degrees: 0^th^ degree, 2^nd^ degree, and 4^th^ degree and obtain a feature vector consisting of three RISH coefficients per shell. For multi-shell acquisitions, we concatenate the RISH feature vectors from each shell. We compute symmetric inverse divergence in line with previous work as follows:

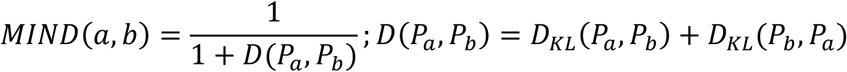

where *a*, *b* signify the pair of regions and *P*_*a*_, *P*_*b*_ their respective probability distributions. We estimate the multivariate KL-divergence via the k-nearest neighbors approach pioneered by Wang et al. [Wang et al., 2009] and used in MIND networks [Sebenius et al., 2023].

### Human Connectome Project Data

We used structural, diffusion, and functional preprocessed data from the S1200 release of the HCP-YA dataset [Van Essen et al., 2013] to create structural and functional connectomes and MSNs, MIND networks, and DSNs. We exclude any subjects with quality control issues due to anatomical anomalies, segmentation and surface errors, temporal head coil instability, and model fitting irregularities [Marcus et al., 2013]. As a result, our analysis comprises 962 subjects, 38 of whom also have retest data. Throughout our analysis, we used the Desikan parcellation [Desikan et al., 2006], consisting of 68 cortical regions (34 per hemisphere) and 14 subcortical regions.

Diffusion data was denoised via Marchenko-Pastur Principal Component Analysis denoising [Veraart et al., 2016] followed by Rician debiasing [Gudbjartsson and Patz, 1995]. Structural T1w MRIs were parcellated via SynthSeg [Billot et al., 2021] and registered to dMRI with boundary-based registration [Greve and Fischl, 2009]. Functional connectivity (FC) was computed via pairwise correlations across time for each pair of regions. Negative correlations were clipped to zero and connectivity matrices were z-transformed, group-averaged, and min-max normalized. Structural connectivity (SC) was computed using MRtrix [Tournier et al., 2019]: multi-shell, multi-tissue response functions were estimated [Jeurissen et al., 2014], spherical deconvolution and intensity normalization were performed, and tractograms with 4 million streamlines were generated. Spherical-deconvolution informed filtering [Smith et al., 2015] was applied, and the streamline count was weighted by inverse node volume and tract length, log-transformed, group-averaged, and min-max normalized. MSNs and MIND networks were computed using previously described methods [Sebenius et al., 2023; Seidlitz et al., 2018].

### MGH-USC Dataset

We used the structural MRI and preprocessed dMRI data from the MGH-USC dataset to establish reproducibility of DSNs and investigate the effect of adding ultra-high b-value shells to their computation. The MGH-USC dataset consists of 35 healthy adults (20-59 years old), of which 32 passed quality-control, with structural (T1w, T2w) and diffusion data acquired on a 3T CONNECTOM scanner, capable of producing gradients up to 300 mT/m. The dMRI acquisition consisted of four shells: 1000 s/mm^2^ (64 directions), 3000 s/mm^2^ (64 directions), 5000 s/mm^2^ (128 directions), and 10000 s/mm^2^ (256 directions) at 1.5 mm isotropic resolution. dMRI was corrected for gradient non-linearity, head motion, and eddy current artifacts. Structural MRI was also corrected for gradient non-linearity. Extensive information on the scanner, acquisition protocols, and preprocessing performed can be found [Fan et al., 2016].

SynthStrip [Hoopes et al., 2022] was used to skull-strip the structural and diffusion images, SynthSeg [Billot et al., 2021] was used to segment and parcellate the T1w structural image, and boundary-based registration (BBR) [Greve and Fischl, 2009; Smith et al., 2004] was used to register the diffusion images to the T1w structural image with the aid of a white matter mask derived from SynthSeg. The SynthSeg cortical parcellation was resampled to the diffusion space via nearest-neighbor resampling and DSN computation proceeded as previously described. We fit two DSNs: one on a limited subset of the dMRI acquisition (only b=1000 s/mm^2^ and b=3000 s/mm^2^) to emulate the b-value range of the HCP-YA dataset and the full acquisition to investigate how DSN characteristics change with the addition of high b-value shells. We applied the same biological fidelity analysis to the DSNs derived from the MGH dataset to ensure fair comparison with those derived from the HCP-YA dataset.

### Reliability Analysis

We evaluate DSNs based on reliability, biological validity, and sensitivity to demographic and cognitive variation. To assess reliability, we compute the test-retest correlation coefficients and the two-way mixed, single measures, absolute agreement intra-class correlation (ICC) over the edge weights. The test-retest correlation coefficients were found by computing the Pearson correlation coefficient over the edge weights for each of the 38 subjects that had test and retest data available. We computed the ICC for each edge by measuring the variation between edge weights using the test-retest portion and used the whole dataset to measure the variation between subjects. Finally, we measured the Pearson correlation coefficient across the edge weights for every pair of subjects as a measure of between subject consistency.

### Biological Validity Analysis

To evaluate biological validity, we measure what percentage of the top 0.1 – 10 % of edges are between regions of the same cytoarchitectural class, as defined by the von Economo and Koskinas atlas [Constantin von Economo et al., 1925; Scholtens et al., 2018]; the same laminar class, as defined by the Mesulam atlas [Mesulam, 1998]; or homologous, connecting corresponding regions across cerebral hemispheres. We compute the assortativity of various micro-architectural maps, such as the sensorimotor-association (SA) axis [Sydnor et al., 2021], cortical thickness [Fischl, 2012], myelination as defined by T1w/T2w ratio [Glasser and Van Essen, 2011], the principal gradient of gene expression [Hawrylycz et al., 2012; Markello et al., 2021], synapse density as measured by SV2A receptor density [Silva-Rudberg et al., 2024], Big Brain structural gradients [Wagstyl et al., 2020], glycolytic index, cerebral metabolic rate of glucose [Vaishnavi et al., 2010], and MEG timescale [Shafiei et al., 2022]. Assortativity was defined as the Pearson correlation between annotations for the weighted connectome [Newman, 2003].

### Age and Cognition Prediction

We trained ridge regression models with tunable ridge penalty using the sklearn package [Pedregosa et al., 2012] to predict age and total, crystallized, and fluid cognition. We evaluated the predictability of the edge weights and the degree centrality for each network via repeated five-fold cross-validation (20 repeats, 100 total folds). Within each fold, the optimal ridge penalty was determined via leave-one-out cross-validation on the training set and predictability was evaluated by measuring the coefficient of determination on the validation set.

### Sex Prediction

We trained logistic regression models with tunable L2 penalty using the sklearn package [Pedregosa et al., 2012] to predict sex. We computed the predictability of the edge weights and the degree centrality for each network via repeated stratified five-fold cross-validation (20 repeats, 100 total folds). Within each fold, the optimal ridge penalty was determined via repeated stratified five-fold cross-validation (4 repeats, 20 total folds) on the training set and predictability was evaluated by measuring the area under the receiver operating curve (AUC-ROC) on the validation set.

### Region Leave-one-out Cross Validation

To determine the importance of each region’s degree centrality to the prediction of age, cognition, and sex, we adopted a region-wise leave-one-out cross validation paradigm. For each fold, one region was removed, the same training and validation protocol took place, and the resulting drop in the coefficient of determination or AUC-ROC was recorded. Regions which recorded a greater drop in performance were surmised to play a greater role in prediction.

### Heritability Analysis

We used the umx package [Bates et al., 2019] to conduct twin-based heritability analysis via structural equation modelling of the ACE model [Maes, 2014], where additive genetic effects, the common environment, and the unique, random environment are considered. We measured heritability, defined as the proportion of the total variance due to additive genetic effects, across the edge weights and degree centralities.

### Macaque Tract Tracing Analysis

MSN and MIND networks were derived using structural MRI (0.3 mm isotropic resolution) from nineteen anesthetized female rhesus macaque monkeys (Macaca mulatta, ages 18.5–22.5 years) from the UC-Davis cohort provided by PRIME-DE [Milham et al., 2018]. The data was pre-processed using the HCP Non-Human Primate pipeline [Hayashi et al., 2021] by Xu et al. [Xu et al., 2020]. Tract tracing correlation coefficients were sourced from the MIND publication [Sebenius et al., 2023]. DSNs were derived from ex-vivo diffusion MRI (0.6 mm isotropic resolution) from six (n=4 male, n=2 female) rhesus macaque monkeys (Macaca mulatta, ages 4.03–15.81 years) consisting of 16 b = 0 s/mm^2^ and 128 b = 4000 s/mm^2^ volumes.

We used four tract-tracing connectomes based on the 91-region Markov M132 parcellation of the left hemisphere: 29 x 29 and 29 x 91 from the original Markov dataset [Markov et al., 2014] and 40 x 40 and 40 x 91 from a recent extension of the Markov dataset [Froudist-Walsh et al., 2021]. We used the log-transformed fraction of labelled neurons (FLNe) as the tract-tracing measure of axonal connectivity. An additional bihemispheric parcellation estimated using structural connectivity measured via dMRI [Shen et al., 2019a] was used to assess agreement over the whole macaque brain. We computed Pearson correlation coefficients for efferent and afferent connections, Fischer-transformed the coefficients, averaged them, and then inverse transformed the value to calculate an aggregate value.

### Statistical Analysis

DSN gradients were computed via degree-normalized Laplacian embedding. We used spin permutations for testing statistical significance with a significance level of 5% [Markello and Misic, 2021]. All multiple hypothesis tests underwent false discovery rate correction. Ridge regression was performed via five-fold repeated nested cross-validation to determine the optimal shrinkage penalty; for each fold, the coefficient of determination was reported on a held-out test dataset.

### Data and Code Availability

Code used to perform DSN analysis can be found at https://github.com/ucsfncl/DSN. We computed MSN, MIND, FC, SC, and DSN connectomes using the Human Connectome Project Young Adult (HCP-YA) dataset (https://db.humanconnectome.org/) [Van Essen et al., 2013]. We sourced gene expression from enigma (https://enigma-toolbox.readthedocs.io/en/latest/index.html) [Larivière et al., 2021], factor maps from a recent publication (https://github.com/LeonDLotter/CTdev/) [Lotter et al., 2024], and other maps from neuromaps (https://netneurolab.github.io/neuromaps/index.html) [Markello et al., 2022]. We made use of the macaque datasets available from PRIME-DE [Van Essen et al., 2012; Milham et al., 2018], using tract tracing data that has been compiled [Froudist-Walsh et al., 2021; Markov et al., 2014] along with the virtual brain data. Cortical surfaces can be accessed at https://balsa.wustl.edu/reference/976nz, tract-tracing data at https://core-nets.orgm, the virtual brain multimodal connectome at https://zenodo.org/record/1471588#.YqBt5S2ca_U, and the F99 template at https://scalablebrainatlas.incf.org/.

## Results

### DSN Estimation and Comparison

We summarize the pipeline for constructing DSNs in Fig. 1. DSNs have a great deal of flexibility and can be computed with varying RISH degree, diffusion weighting, spatial and angular resolution, and parcellation. In our analysis, we consider DSNs computed using multi-shell acquisitions as well as DSNs computed using a faster, more clinically feasible single-shell b=1000 s/mm^2^ acquisitions (30 directions) on the Desikan-Killiany (DK) parcellation [Desikan et al., 2006] with 1.25 mm isotropic resolution. We compare these multi-shell and single-shell DSNs to other similarity networks, namely MSNs and MIND networks, as well as SC from dMRI tractography and functional connectivity FC from rs-fMRI seed-based analysis [Larivière et al., 2021] (Figs. 2, 3). Like these other forms of brain network analysis, the DSNs are also largely bilaterally symmetric. Most of our analysis is conducted using the Human Connectome Project Young Adult (HCP-YA) dataset [Van Essen et al., 2013]. Additional detail is provided in the Methods.

**Figure 1:**
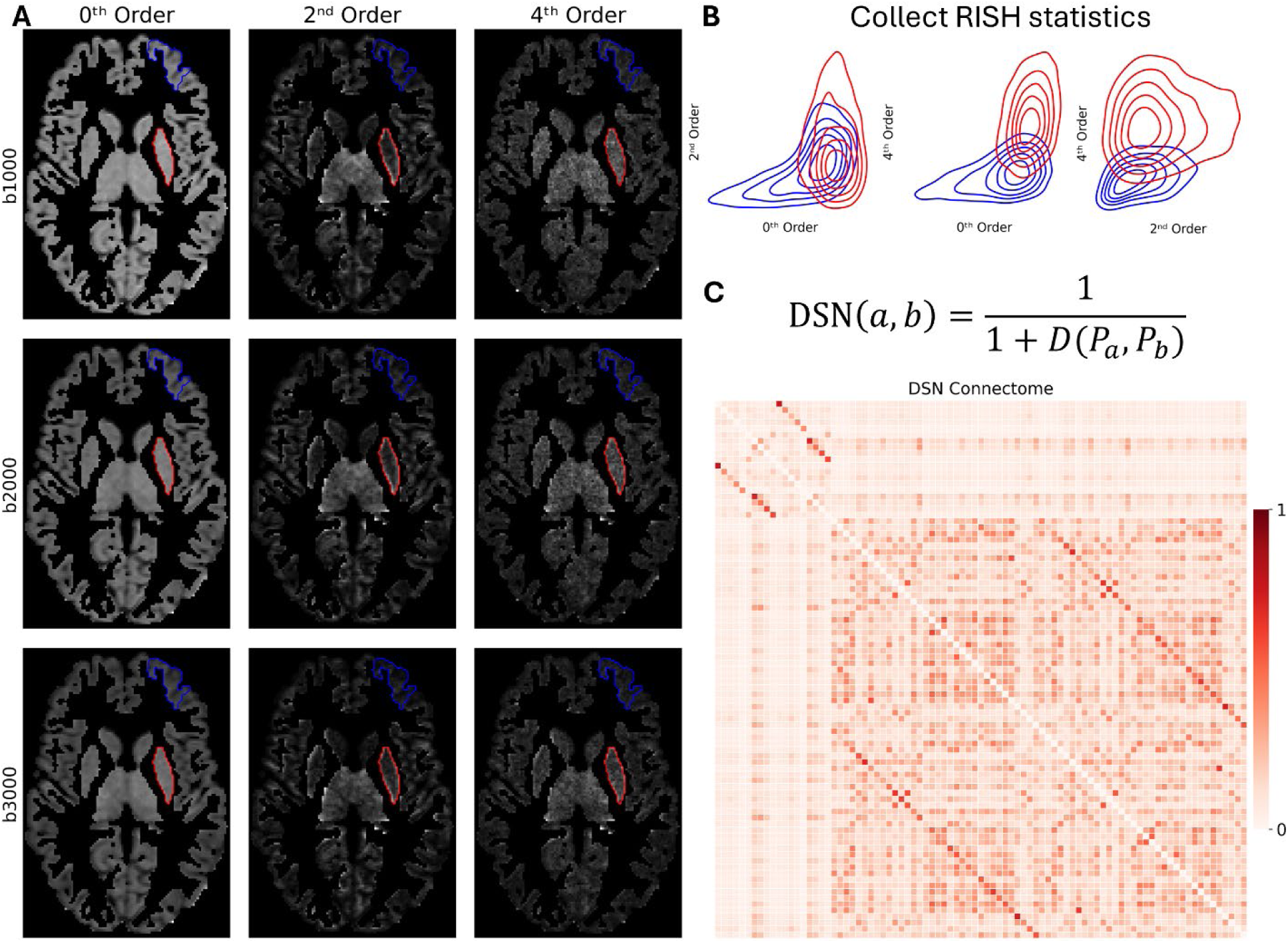
Computing Diffusion Similarity Networks. (A) The 0^th^, 2^nd^, and 4^th^ degree rotationally invariant spherical harmonics (RISH) maps for the b=1000, 2000, 3000 s/mm^2^ shells across the gray matter of an example subject. Diffusion similarity estimation between two example regions, the left pars opercularis (blue) and left putamen (red), is shown. (B) The RISH statistics are collected for both regions (only b=1000 s/mm^2^ is shown) and (C) used to compute the Diffusion Similarity Network (DSN) connectome via the inverse symmetric Kullback-Leibler (KL) divergence.

**Figure 2:**
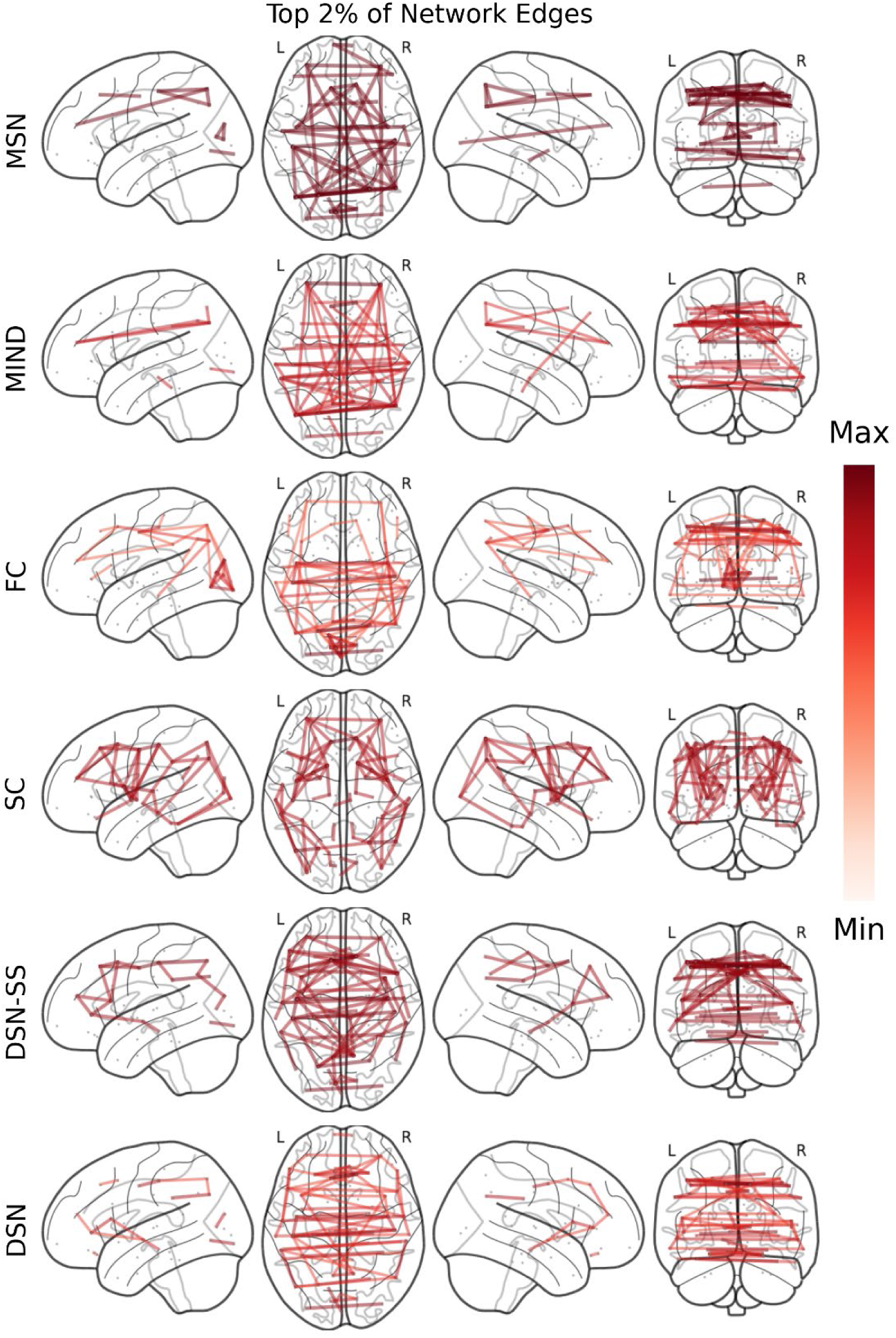
Visual Comparison of Network Edges. The strongest 2% of edge weights of the following group-averaged networks (n=962): morphometric similarity networks (MSNs), morphometric inverse divergence (MIND) networks, functional connectivity (FC), structural connectivity (SC), and diffusion similarity networks computed from single shell (DSN-SS) and multi-shell (DSN) acquisitions.

**Figure 3:**
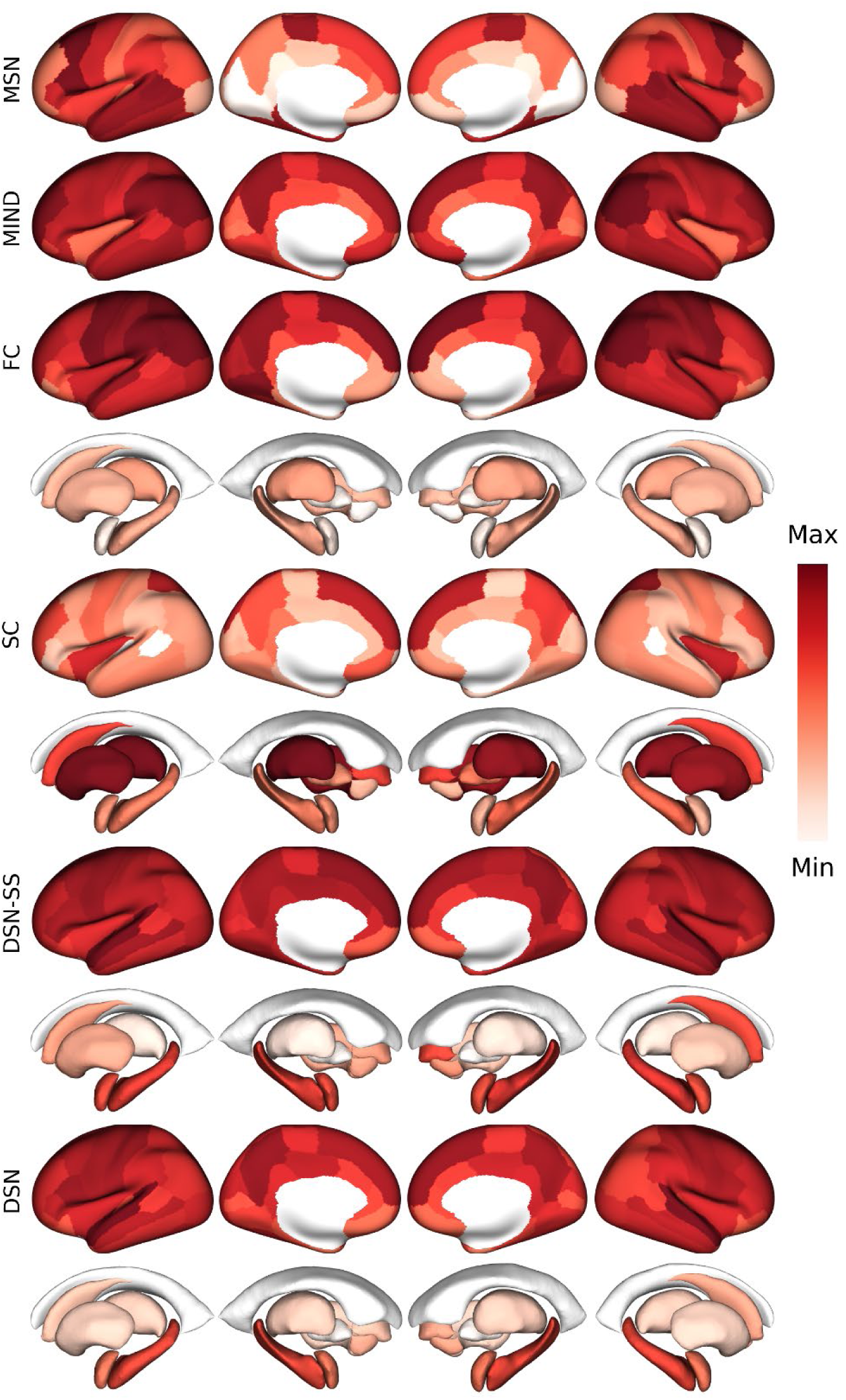
Visual Comparison of Network Degree. The network degree of the following group-averaged networks (n=962): morphometric similarity networks (MSNs), morphometric inverse divergence (MIND) networks, functional connectivity (FC), structural connectivity (SC), and diffusion similarity networks computed from single shell (DSN-SS) and multi-shell (DSN) acquisitions.

### Network Reliability

We first investigated the reliability of DSNs computed using single-shell and multi-shell acquisitions by recording the Pearson correlation coefficient between every pair of subjects (Fig. 4a) and between test-retest sessions (Fig. 4b) as well as the intra-class correlation coefficient (ICC) (Fig. 4c) across the edge weights. DSNs achieved significantly greater between-subject correlations than MSN (t=524, p<1e-12), FC (t=141, p<1e-12), and SC (t=296, p<1e-12) using two-sided paired t-tests. Multi-shell DSNs achieved excellent test-retest correlation coefficients (r=0.92, CI=[0.86, 0.96]) and ICCs (r=0.86, CI=[0.74, 0.94]). Despite their worse performance, single-shell DSNs still achieved good reliability with high test-retest correlation coefficients (r=0.83, CI=[0.75, 0.91]) and ICCs (r=0.83, CI=[0.71, 0.92]).

**Figure 4:**
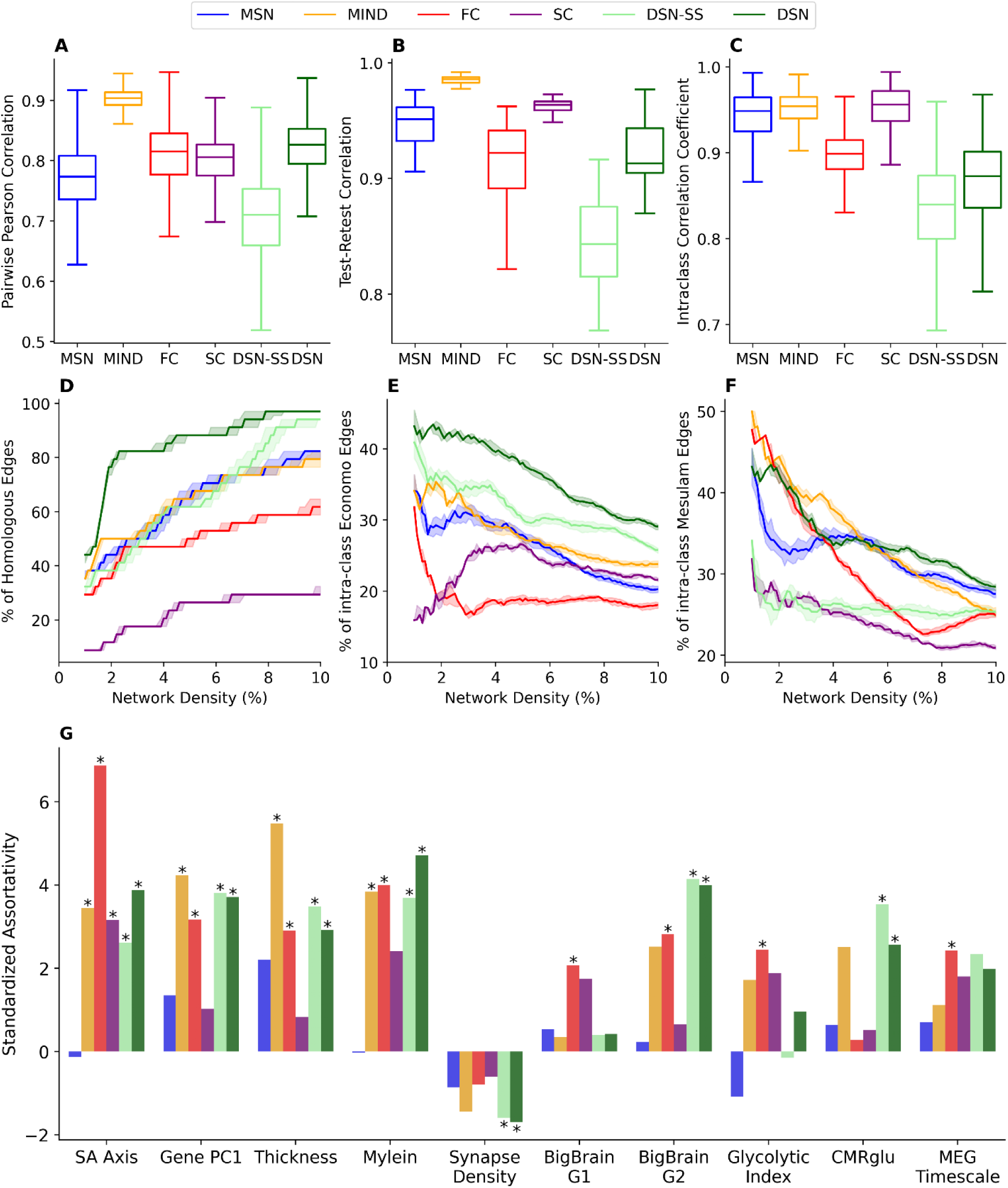
Comparison of Network Connectomes. (A) between subject Pearson correlation, (B) test-retest Pearson correlation, and (C) intra-class correlation coefficients as well as (D) the percentage of connections between regions that are homologous, (E) within the same cytoarchitectural von-Economo and Koskinas class, and (F) within the same Mesulam class over varying network densities (1% - 10%) using group-averaged networks (n=968). The shaded regions (D-F) represent the 95% confidence interval estimated by subject bootstrapping. (G) The network assortativity standardized against the null-distribution, found via spin permutations, of the sensorimotor-association (SA) axis, thickness, myelin, synapse density measured using synaptic vesicle glycoprotein 2A (SV2A) receptor binding, BigBrain histological gradients (G1 & G2), glycolytic index (GI), cerebral metabolic rate of glucose consumption (CMRglu), and magnetoencephalography timescale. Asterisks indicate significance of assortativity, determined by two-sided spin tests. Between subject Pearson correlation was found by recording the correlations between every pair of subjects over the edge weights (n = 968). Test-retest Pearson correlation was computed by recording the Pearson correlation between the edge weights computed from the test and retest sessions (n=38). We use the two-way mixed, single measures, absolute agreement inter-class correlation and measure the variation between edge weights using the test-retest portion and variation between subjects using the whole dataset. Statistics were acquired for morphometric similarity networks (MSNs), morphometric inverse divergence (MIND) networks, functional connectivity (FC) networks, structural connectivity (SC) networks, and diffusion similarity networks computed using a single shell at b = 1000 s/mm^2^ with 30 directions (DSN-SS) and a multi-shell acquisition (DSN). Networks were constructed using the 68 region Desikan parcellation with only cortical regions and cortico-cortical connections considered.

### Biological Validity

Next, we evaluated the extent to which single-shell and multi-shell DSNs exhibited the following three principles of neural organization: bilateral symmetry, cytoarchitecture, and cortical laminar differentiation at varying network density thresholds from 1% to 10%. Compared to all other networks, multi-shell and single-shell DSNs had a significantly greater percentage of connections that were between homologous inter-hemispheric regions (Fig. 4d) and within the same von Economo-Koskinas cytoarchitectonic class (Fig. 4e) across all thresholds with two-sided bootstrap tests with false discovery rate correction. Multi-shell DSNs had a significantly greater percentage of connections between regions of the same Mesulam class than MIND (5.8% - 10%) and FC (2.5% - 10%), MSNs and SC for all thresholds (Fig. 4f). In comparison, single-shell DSNs had a much lower percentage of intra-class Mesulam connections, only greater than that of SC (3.8% - 10%) and FC (6.5% - 9.3%).

We replicated our biological validity analysis on the MGH-USC dataset, focusing onDSNs computed using a limited subset of the dMRI acquisition (only b=1000 s/mm^2^ and b=3000 s/mm^2^) and a full acquisition (up to b=10000 s/mm^2^). The DSNs computed using the MGH-USC achieved similar results as those computed using the HCP-YA dataset. The inclusion of ultra-high b-value shells (b=5000 s/mm^2^ and b=10000 s/mm^2^) led to a significantly greater percentage of homologous connections (6.6% - 10%) captured, but no such increase in intra-class von Economo-Koskinas and intra-class Mesulam connections (Fig. S1).

In addition, we investigated the ability of multi-shell and single-shell DSNs to capture various macroscale, microarchitectural, electrophysiological and metabolic characteristics of the cortex, such as the SA axis, myelination measured via T1w/T2w ratio, thickness, synaptic density measured via synaptic vesicle glycoprotein A (SV2A) receptor binding density, histological gradients (BigBrain G1 & G2), glycolytic index (GI), cerebral metabolic rate of glucose consumption (CMRglu), and magnetoencephalography (MEG) timescale via network assortativity (Fig. 4g). We found multi-shell DSNs to have significant assortativity with the SA axis (p=0.003), thickness (p=0.012), myelin (p=0.0027), synaptic density (p=0.03), BigBrain G2 (p=0.0031), and CMRglu (p=0.041) following two-sided spin permutation tests with false discovery rate (FDR) correction. Single-shell DSNs did not lag far behind with statistically significant assortativity with SA-axis (p=0.043), thickness (p=0.01), myelin (p=0.01), SV2A (p=0.007), BigBrain G2 (p=0.006), and CMRglu (p=0.015). In comparison, MSNs did not have any significant assortativity; MIND had significant assortativity with SA-axis (p=0.015), thickness (p=0.002), and myelin (p=0.01); FC had significant assortativity with SA-axis(p=0.0018), thickness (p=0.03), myelin (p=0.014), BigBrain G1 (p=0.039), BigBrain G2 (p=0.021), GI (p=0.033), and MEG timescale (p=0.034); SC had significant assortativity with only the SA axis (p=0.021).

Then, we explored the ability of DSN to capture homophily, or the tendency of structurally similar regions to be anatomically connected via white matter tracts, in comparison to other similarity measures. We found DSN, along with all networks, to have significantly greater similarity between structurally connected regions (t=13.7, p=1e-38) using two-sided independent t-tests with unequal variances and to be significantly correlated with structural connectivity as measured via dMRI tractography over cortico-cortical connections in living human brains (r=0.37, p=0.002) (Fig. S2). We did not find DSN to have significantly greater similarity for structurally connected regions or significant correlation with structural connectivity over subcortico-cortical or subcortico-subcortical connections.

Lastly, we extended validation from human in-vivo dMRI tractography to an ex-vivo macaque model with retrograde tract tracing, specifically the fraction of labelled neurons (FLNe), as a gold standard measure of axonal connectivity. We analyzed a single shell dMRI dataset (b=4000 s/mm^2^) with 0.5 mm isotropic resolution comprised of six ex-vivo macaque brains. We made use of the work of Seidlitz et al. [Seidlitz et al., 2018] and Sebenius et al. [Sebenius et al., 2023], considering five connectomes [Froudist-Walsh et al., 2021; Markov et al., 2014; Shen et al., 2019b] based on two cortical parcellations [Markov et al., 2014; Shen et al., 2019b], and found DSN to be better correlated with tract-tracing measures than MSN, but worse so than MIND (Fig. S3).

### Diffusion Similarity Network Gradients

To gain deeper insight into the structural organization of DSNs, we computed gradients of the group-averaged DSN via normalized Laplacian embedding. The first DSN gradient (G1) displayed a top-down neocortical-subcortical hierarchy (Fig. 5a), whereas the second DSN gradient (G2) (Fig. 5b) was significantly correlated with the SA-axis (r=0.78, p=0.0028) (Fig. 5c) and was stratified across the SA & sensory-fugal axes and distinguished polar cortex (Fig. 5d). DSN G2 was significantly correlated with dopaminergic/glutaminergic receptor densities (FDOPA-DAT-D1-NMDA: r= 0.64, p=0.0285), flow & metabolism (CBF-CMRglu: r=-0.61, p=0.0285), immunity (COX1: r=-0.66, p=0.0154), general neuronal expression (Ex7-In4-Ex5-Ex4-In1: r=0.84, p=0.0071), microglia/oligodendrocyte progenitor (Micro-OPC: r=0.65, p=0.0389), and inhibitory/astrocyte (In3-In2-Astro: r= 0.73, p=0.0285) factors (Fig. 5e). DSN G2 was also correlated with cortical thickness (r=0.77, 0.0048) and myelin (r=-0.71, p=0.0066); gene expression (r=0.85, p=0.004), BigBrain G2 (r=-0.85, p=0.0048), and FC (r=0.68, p=0.006) gradients; MEG power bands for delta (2-4 Hz: r=0.84, p=0.0368) and high gamma (60-90 Hz: r=0.78, p=0.0204) bands as well as MEG timescale (r=0.74, p=0.0368) (Fig. 5f). Significance was assessed via two-sided spin tests with false discovery rate correction.

**Figure 5:**
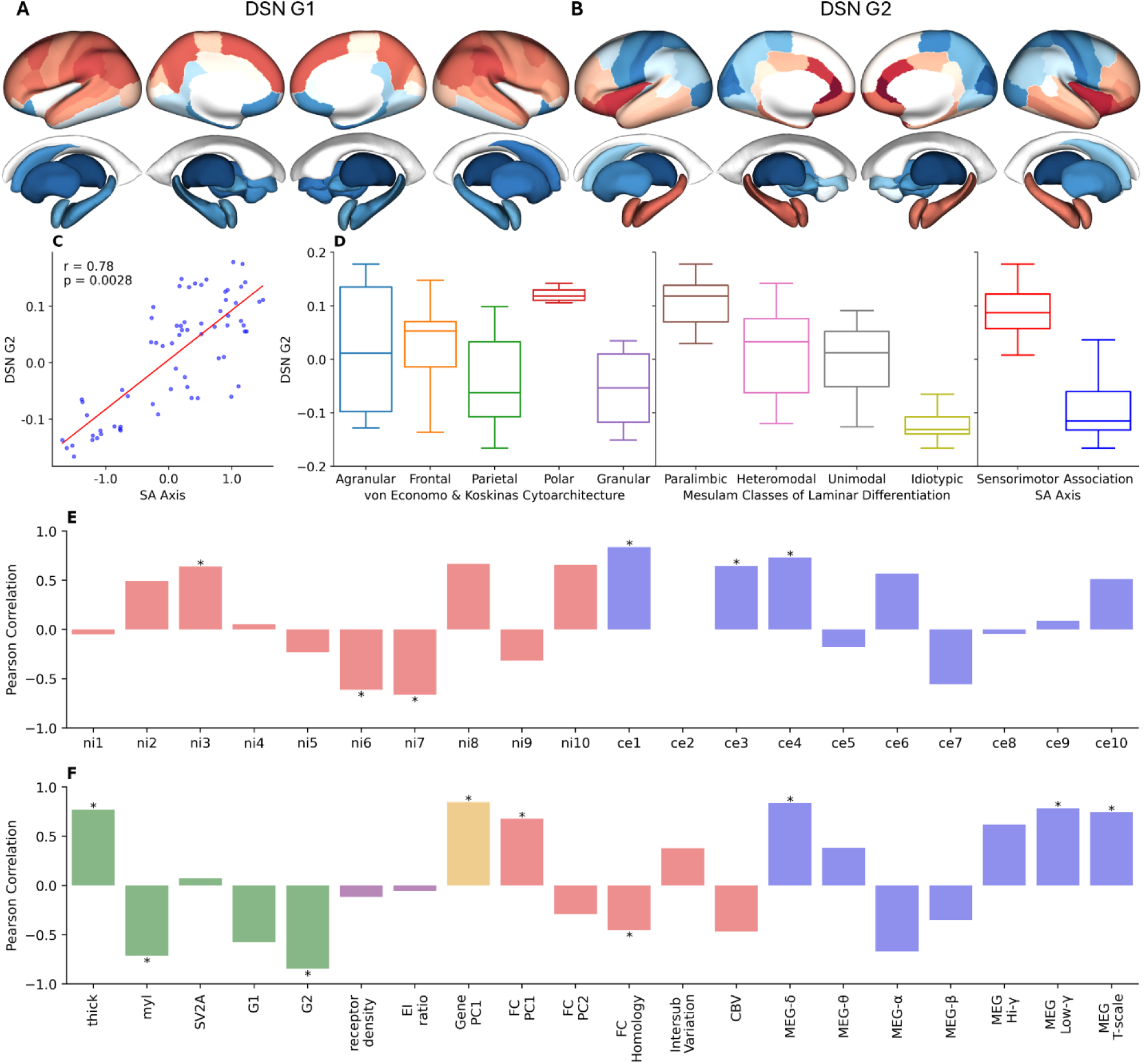
The Gradients of Diffusion Similarity Networks. (A) The first & (B) second gradients (G1 & G2) derived from degree-normalized Laplacian embedding of the group-averaged Diffusion Similarity Network (DSN). (C) DSN G2 aligns with the sensorimotor-association (SA) axis (r=0.78, p=0.0028), (D) distinguishes polar cortex among the von Economo classes, is stratified across the sensory-fugal axis of Mesulam, and differentiates sensorimotor from association cortex. DSN G1 shows hierarchical top-down organization from neocortex to paleocortex/archicortex to subcortical gray matter. (E) Univariate correlations were taken between the second gradient and ten neuronal and cellular factors respectively. DSN G2 was significantly correlated with dopaminergic/glutaminergic receptor densities (ni3-FDOPA-DAT-D1-NMDA: r= 0.64, p=0.0285), flow & metabolism (ni6-CBF-CMRglu: r=-0.61, p=0.0285), immunity (ni7-COX1: r=-0.66, p=0.0154), general neuronal expression (ce1-Ex7-In4-Ex5-Ex4-In1: r=0.84, p=0.0071), microglia/oligodendrocyte progenitor (ce3-Micro-OPC: r=0.65, p=0.0389), and inhibitory/astrocyte (ce4-In3-In2-Astro: r= 0.73, p=0.0285) factors. We investigated the relationship between DSN G2 and cortical annotations further: DSN G2 was significantly correlated with cortical thickness (r=0.77, 0.0048) and myelin (r=-0.71, p=0.0066); gene expression (r=0.85, p=0.004), BigBrain G2 (r=- 0.85, p=0.0048), and FC (r=0.68, p=0.006) gradients; MEG power bands for delta (2-4 Hz: r=0.84, p=0.0368) and high gamma (60-90 Hz: r=0.78, p=0.0204) bands as well as MEG timescale (r=0.74, p=0.0368). Significance was assessed via two-sided spin tests with false discovery rate correction.

### Sensitivity to Age, Cognition, and Sex

After evaluating the reliability and neurobiological validity of DSNs, we gauged their sensitivity to detect relevant inter-individual variation due to age, cognition, and sex in the HCP-YA cohort (n=968). We trained ridge regression models via repeated five-fold cross validation (20 repeats, 100 folds), using the training set for tuning the ridge penalty via leave-one-out cross validation and the validation set to compute the coefficient of determination for age and fluid, crystallized, and total cognition using edge weight and degree centrality features from MSN, MIND, FC, SC, and DSNs computed from a b=1000s/mm^2^ shell (30 directions) and a multi-shell acquisition. Logistic regression models were trained via repeated stratified five-fold cross validation (20 repeats, 100 folds), using the training set for tuning the L2 penalty via repeated stratified cross validation (4 repeats, 20 folds) and the validation set to compute the area under the receiver operator curve (AUC-ROC) for sex classification using edge weight and node centrality features from the same networks. All models were trained and tested on the same folds. We evaluated the differential performance of multi-shell and single-shell DSNs via one-sided paired t-tests with false discovery rate correction.

Edge weight regression models trained using DSNs significantly outperformed MSNs and MIND on the prediction of fluid cognition, MSNs, MIND, and SC on the prediction of total cognition, and all other networks on the prediction of crystallized cognition and age (Fig. 6a). Single-shell DSNs performed worse than their multi-shell counterparts, but still significantly outperformed the other networks, apart from FC on crystalized cognition. Degree centrality ridge regression models trained using DSNs significantly outperformed all other networks for prediction of fluid, crystalized, and total cognition and age (Fig. 6b). Once again, single-shell DSNs performed worse than their multi-shell counterparts, especially for the prediction of fluid cognition, where they only outperformed FC, but outperformed all other networks on all other regression targets. Logistic regression models trained using multi-shell DSN edge weight and node centrality achieved significantly higher AUC-ROC than all other networks (Fig. 6c, 6d). Single-shell DSNs performed comparably but achieved slightly lower AUC-ROC than SC using edge weights (Tables S1, S2).

**Figure 6:**
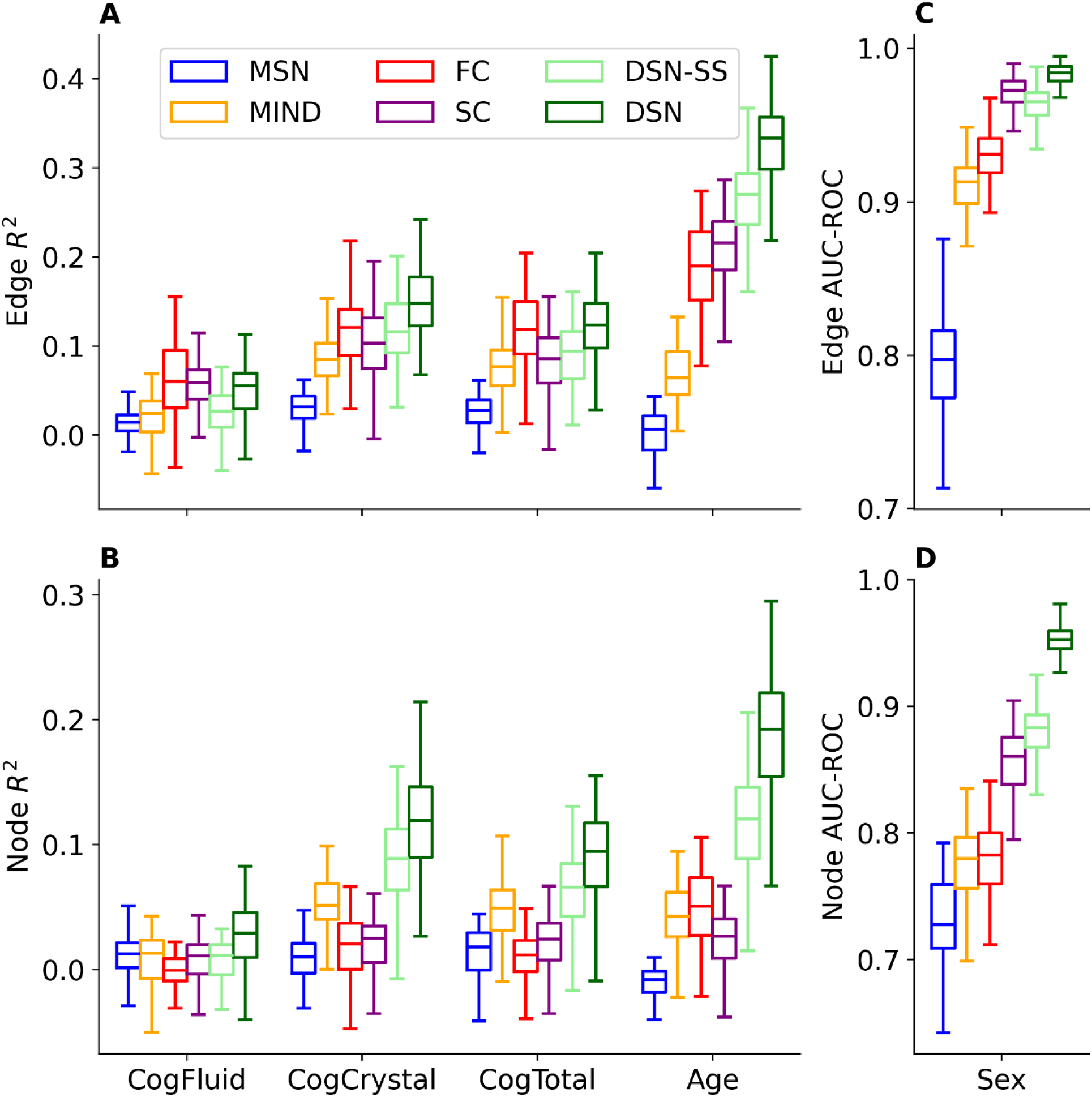
Prediction of Age, Cognition, and Sex with Network Connectomes. The coefficient of determination (R^2^) for the prediction of fluid, crystallized, and total cognition along with age using (A) edge weights and (B) node centrality as well as the area under the receiver-operator curve (AUC-ROC) for classification of sex using (C) edge weights and (D) node centrality. For regression tasks, ridge regression models were trained with repeated five-fold cross-validation (20 repeats, 100 folds); the optimal ridge penalty was found via leave-one-out cross-validation on the training set and performance was evaluated on the validation set. For classification of sex, logistic regression models were trained with repeated stratified five-fold cross-validation (20 repeats, 100 folds); the optimal L2 penalty was found via repeated stratified five-fold cross-validation (4 repeats, 20 folds) on the training set and performance was evaluated on the validation set.

To determine which regions contributed most to age and sex prediction, we performed the same cross-validation schemes with one region left out and reported the corresponding decrease in the coefficient of determination and AUC-ROC respectively (Fig. 7). We found most of the sensitivity of DSNs to age comes from perirolandic and mesial temporal regions, and their sensitivity to sex to stem from lateral parieto-occipital areas especially within the right hemisphere. MIND showed similar results in the neocortex for age and sex but could not be applied to allocortical regions of mesial temporal lobes such as the amygdalohippocampal complex. The areas identified by DSN were also those that had the largest univariate correlation coefficients for age and effect size with respect to sex (Fig. S4). In a similar vein, the top 1% of DSN edge correlation coefficients with respect to age consisted of positive correlations between primary visual regions, such as the pericalcarine, and secondary visual regions and negative correlations between the thalamus and motor regions. The top 1% of DSN effect sizes with respect to sex consisted of connections between the left hippocampus/hippocampal gyrus and visual processing areas, such as the inferior-parietal and cuneus (Fig. S5). Correlation of these correlation coefficients and effect sizes between networks indicated greater agreement among structural similarity networks (MSN, MIND, and DSN) as well as SC for age and less agreement over sex, with only MSN and DSN-SS & DSN showing significant agreement (Fig. S6).

**Figure 7:**
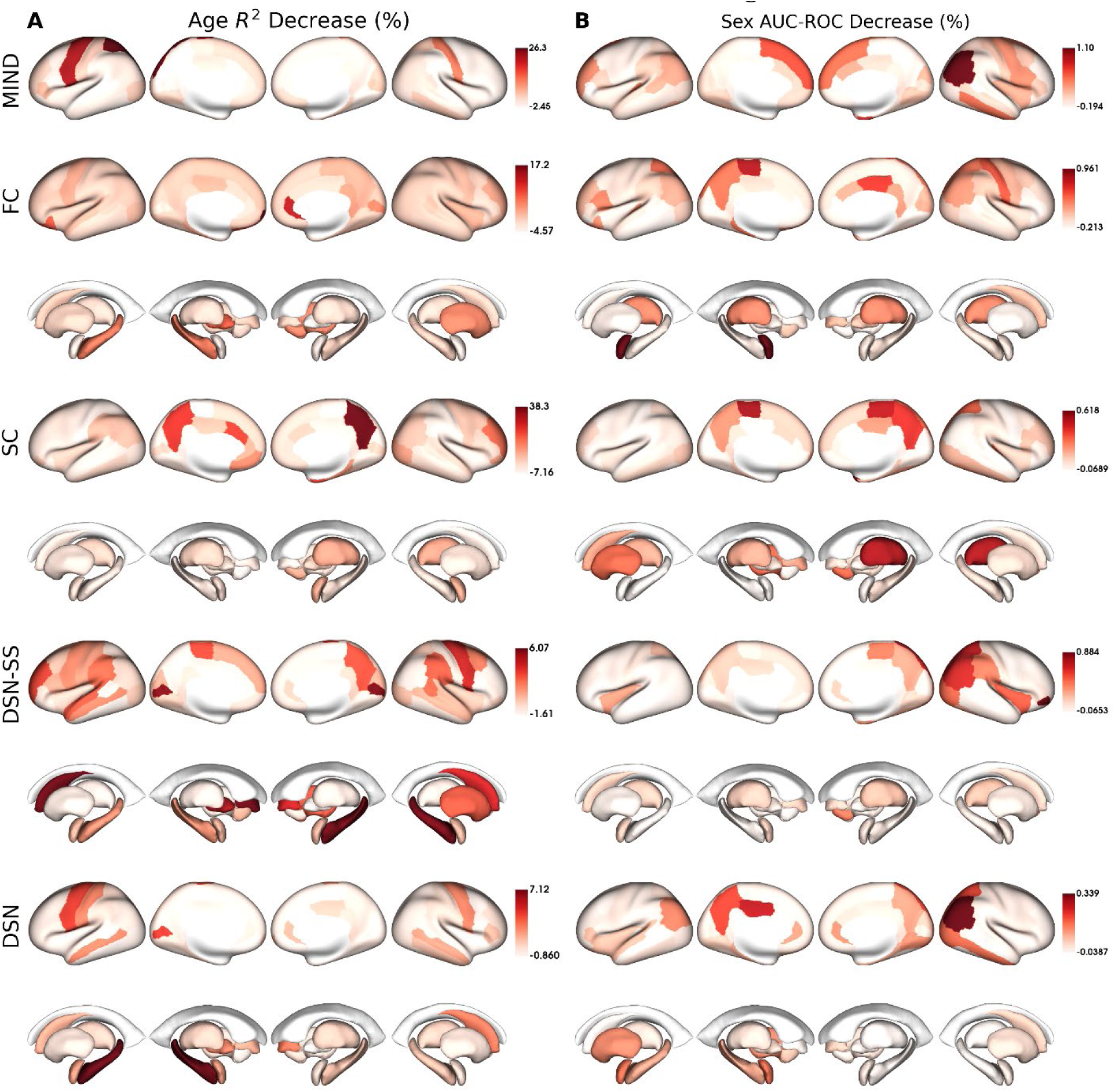
Region-wise Leave-one-out Cross Validation for Age and Sex Prediction. The mean percentage decrease in (A) the coefficient of determination for age regression and (B) the area under the receiver-operator curve (AUC-ROC) for sex classification when each region is left out of the ridge/logistic regression models using network degree. There was no change to the cross-validation scheme, or the folds compared to the model using all regions.

### Similarity to other Brain Networks

We next evaluated the similarity of single-shell and multi-shell group-averaged DSNs to other group-averaged brain networks including gene expression networks by computing inter-network Pearson correlation coefficients across the edge weights and degree centralities. We found DSNs to be significantly correlated with gene expression (r=0.48, p=0.0008), MSN (r=0.37, p=0.002), MIND (r=0.55, p=0.001), FC (r=0.56, p=0.0006), and SC (r=0.37, p=0.002) across their cortico-cortical edge weights and MIND (r=0.58, p=0.004), FC (r=0.7, p=0.004), and SC (r=0.37, p=0.028) across their degree centralities following two-sided spin permutation tests with (FDR) correction (Fig. 8). Single-shell DSNs followed much of these patterns, except they were not significantly correlated with SC degree, but were significantly correlated with gene expression degree. DSNs did not show any statistically significant correspondence for subcortico-cortical connections but were significantly correlated with FC edge weights across the subcortico-subcortical connections (r=0.52, p=0.008) (Fig. S7). In addition to direct edge and node-wise correlations, we also assessed the distance between the subspaces spanned by the top ten Laplacian eigenmodes[Belkin and Niyogi, 2001] for each network via the Grassmann distance [Ye and Lim, 2016] (Fig. S8). We found the DSN eigenspace to be more closely aligned with gene expression, FC, and SC eigenspaces than the MIND or MSN eigenspaces.

**Figure 8:**
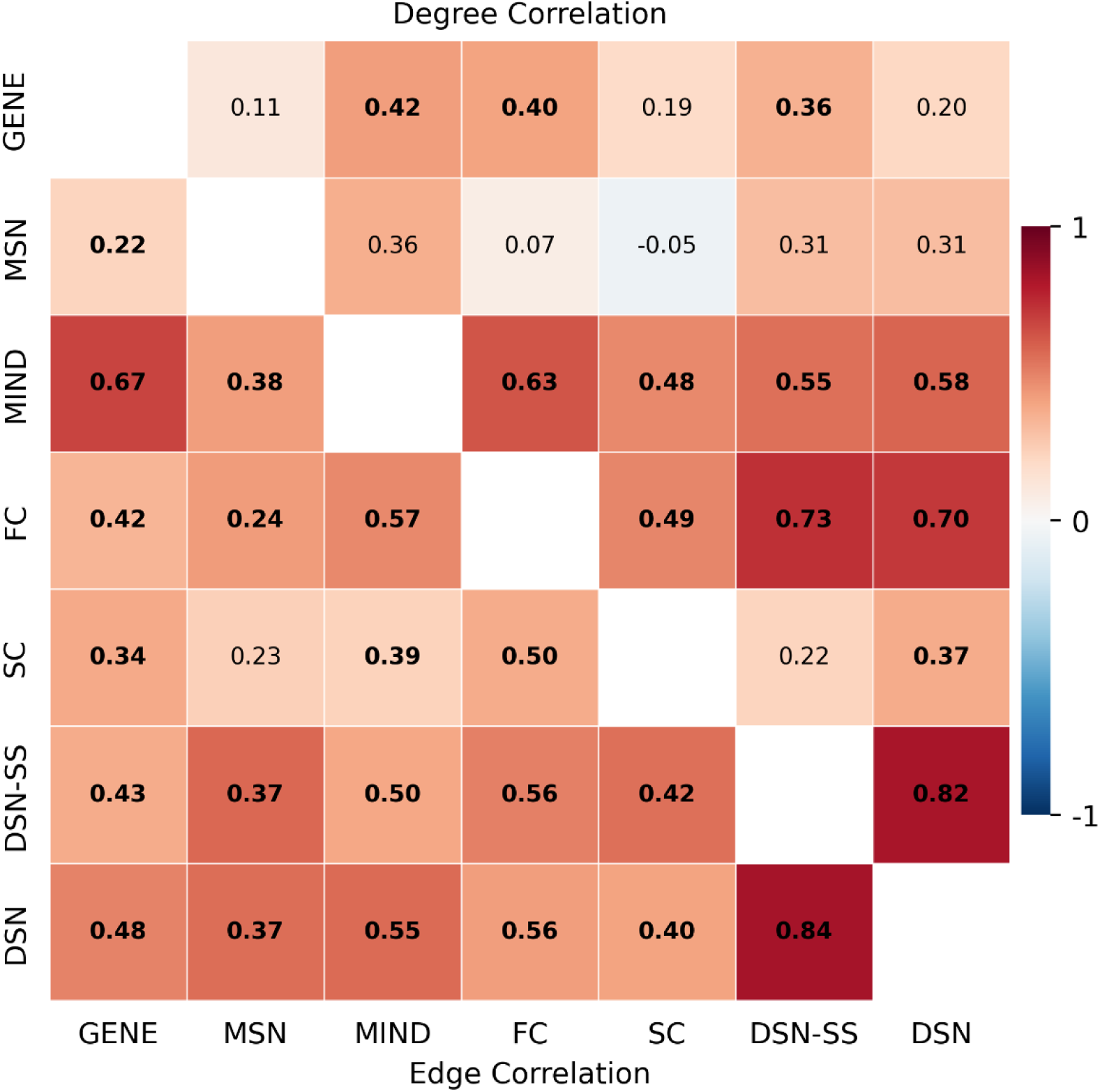
Mapping Similarity between Network Connectomes. The Pearson correlation coefficient between group-averaged network edge weights (*bottom/left*) and node centrality (*top/right*) for cortico-cortical connections. The MIND edge weights better capture gene expression networks, while DSN edge weights better capture structural connectivity. DSN node centrality is better at capturing functional nodal degree, whereas MIND node centrality is better correlated with structural node centrality and gene expression node centrality. Single-shell DSNs (DSN-SS) have slightly lower, but comparable correlations with other networks compared to DSNs. Bolded values indicate statistical significance following two-sided spin tests with false discovery rate correction (computed across all network pairs).

### Edge and Node Heritability

We evaluated heritability over edge weight and node centrality statistics via ACE modelling [Maes, 2014] of monozygotic and dizygotic twin pairs (n=84 dizygotic, 138 monozygotic), considering additive genetic (A), common environment (C), and unique, random environment (E) effects (Figs. S9, S10). We defined heritability as the proportion of the variance that could be explained by additive genetic effects, corrected for age, sex, and estimated intracranial volume.

DSNs were significantly more heritable than MSNs (t=12.1, p=1.3e-32) and MIND networks (t=8.6, p=2.7e-17) across cortico-cortical edges, FC (t=6.68, p=8.1e-11) and SC (t=6.07, p=2.9e-9) across subcortico-cortical edges, FC across subcortico-subcortical edges (t=2.37, p=3.9e-2), and MSNs (t=4.15, p=1.9e-4) and MIND networks (t=3.39, p=1.6e-3) along cortical degree centralities (Fig. 9a, 9b). We evaluated the significance of differential heritability under paired two-sided t-tests with FDR correction. We found DSN heritability to be greater for edges that were structurally wired for cortico-cortical (t=1.9, p=0.058) and subcortico-cortical (t=2.44, p=0.015) using two-sided independent t-tests with unequal variances, mirroring results seen for other networks (Fig. S11).

**Figure 9:**
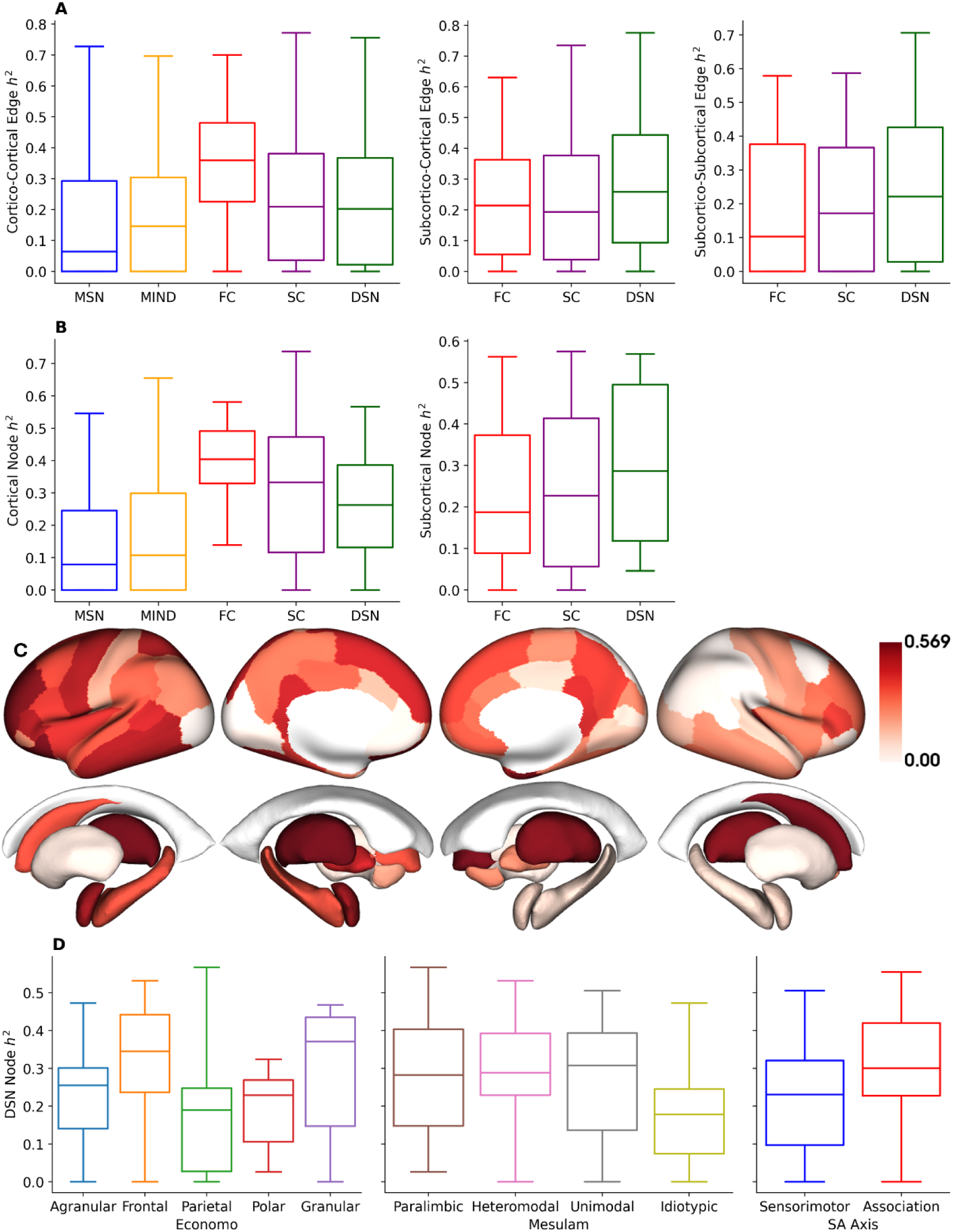
Estimating the Heritability of Network Connectomes. (A) The edge weight heritability (h^2^) across cortico-cortical, subcortico-cortical, and subcortico-subcortical connections and (B) degree heritability across cortical and subcortical nodes. DSNs had significantly higher heritability than MSNs (t=12.1, p=1.3e-32) and MIND networks (t=8.6, p=2.7e-17) across cortico-cortical edges, FC (t=6.68, p=8.1e-11) and SC (t=6.07, p=2.9e-9) across subcortico-cortical edges, FC across subcortico-subcortical edges (t=2.37, p=3.9e-2), and MSNs (t=4.15, p=1.9e-4) and MIND networks (t=3.39, p=1.6e-3) along cortical degree. (C) DSN degree heritability stratified across (D) von Economo & Koskinas cytoarchitectonic classes, Mesulam classes of laminar differentiation, and the sensorimotor-association (SA) axis. DSN had its highest heritability in heteromodal association areas and lowest in idiotypic sensory regions.

DSN degree was most heritable in heteromodal association areas and least heritable in idiotypic sensory regions (Fig. 9c, 9d). DSN and SC degree heritability showed similar stratification across the cortex, whereas FC degree heritability was higher in associations areas and lower in sensorimotor areas (Fig. S12). Apart from MSN and MIND, no other networks had statistically significant associations between their heritability over edge weights or degree centralities for cortical or subcortical regions (Fig. S13). DSN edge weight was negatively correlated with heritability across cortico-cortical (r=-0.11, p=0.16), subcortico-cortical (r=-0.2, p=0.057), and subcortico-subcortical connections (r=-0.39, p=0.001) using two-sided spin tests, whereas SC and FC edge weights were positively correlated with their corresponding heritability (Fig. S14). We found no significant relationship between DSN node heritability and node weight, although FC and MIND had significant positive associations (Fig. S15).

## Discussion

We introduce DSNs as a way of distilling the rich information content of dMRI into an interpretable mapping of the structural organization of individual living human brains and show DSNs to perform well in comparison to other brain similarity networks and connectomes over a wide range of benchmarking tasks.

DSNs exhibited a stronger coupling with cortical cytoarchitecture, laminar classes, and micro-architecture than MSNs and MIND, more faithfully reflecting fundamental organizational principles of the cortex [Constantin von Economo et al., 1925; Mesulam, 1998; Scholtens et al., 2018]. This was corroborated by further investigation in an independent ultra-high b-value dataset. Moreover, DSNs displayed heightened sensitivity to interindividual variation due to age, sex, and cognition, outperforming not only MSN and MIND, but also conventional structural and functional connectomes in many cases, highlighting their potential as sensitive biomarkers in studies of normative and atypical brain trajectories [Rutherford et al., 2023]. Furthermore, we observed DSNs to have greater heritability than MSNs and MIND, indicating a robust genetic component to DSNs and positioning them as potential tools for genetic investigations of brain structure [Yao et al., 2023].

At a methodological level, the enhanced sensitivity of DSNs can be attributed to its usage of dMRI, specifically RISH features, as opposed to MIND and MSNs which use morphometric features from structural T1w MRI. Unlike morphometry, diffusion MRI directly interrogates microstructure by measuring the diffusion of water in neuronal tissues over micron spatial scales. Previous literature has found high-resolution ex-vivo maps of DTI and MAP-MRI parameters to reveal laminar substructures [Avram et al., 2022] and show correlation with histological markers of cytoarchitecture [Wang et al., 2020] and in-vivo human dMRI microstructure to organize along Mesulam’s hierarchy of laminar differentiation and distinguish von Economo and Koskinas cytoarchitectonic classes [Sadikov et al., 2024]. Similarly, the greater consistency and reliability of MIND can be attributed to the high consistency and reliability of morphometry.

The biological validity of DSNs is further corroborated by their robust associations with gene expression networks and structural and functional connectivity. In addition, DSN gradients show clear organizational patterns that aid its interpretability: the first gradient reflects top-down organization extending from the neocortex to paleocortex/archicortex to subcortical gray matter and the second is aligned with the sensory-fugal and sensorimotor-association axes. The significant associations of the second DSN gradient with neuronal oscillatory dynamics, metabolism, neuroimmunity, and dopaminergic and glutamatergic neurotransmitter receptor densities suggest that DSNs reflect biologically meaningful variations in regional microstructure and their neurobiological correlates.

DSNs demonstrated sex differences that corresponded to known sexual dimorphisms of the human cerebral cortex, most prominently in the right greater than left parietooccipital regions which have been related to visuospatial functions such as mental rotation [Hugdahl et al., 2006; Jordan, 2002]. These cortical sex differences were also apparent from MIND, but not to as great an extent as from DSNs. DSNs also highlighted the known strong association of medial temporal lobe structures with aging, particularly the hippocampi [Raz et al., 2004]. These allocortical structures cannot be investigated with morphometric similarity methods such as MSNs or MIND.

Apart from the 29 x 29 connectome, DSN had worse agreement with tract tracing measures of axonal connectivity than MIND. One potential reason could be the lower dataset size and quality of the dMRI data. The dMRI dataset consisted of fewer brains (n=6) as opposed to (n=19) and worse spatial resolution of 0.5 mm vs. 0.3 mm resolution for structural MRI. In addition, the strongly reduced diffusivity and T2 of fixed tissue requires significantly greater diffusion weighting for equivalent contrast leading to greater distortion artifacts, while the lower proton density results in worse signal-to-noise ratio in comparison to in-vivo acquisitions [Roebroeck et al., 2019]. Despite these shortcomings in ex vivo imaging, DSN had better correlation with tract tracing than MSN.

Using DSNs, we confirmed homophily over cortico-cortical connections, observing greater similarity between regions that were structurally connected and significant positive correlations with structural connectivity. However, these observations did not hold for subcortico-cortical and subcortico-subcortical connections, suggesting a lack of homophily for these connections. Future work will be needed to corroborate these findings.

In this preliminary investigation of dMRI as a source of structural similarity, we have focused solely on RISH features. However, metrics derived from signal representations, such as DTI [Basser et al., 1994; Pierpaoli et al., 1996], DKI [Jensen et al., 2005], and MAP-MRI [Özarslan et al., 2013], or microstructural models, such as NODDI [Zhang et al., 2012] or NEXI [Jelescu et al., 2022], could prove useful in conjunction with morphometric features and T1w/T2w ratio. We saw limited utility in increasing RISH beyond 4^th^ order or acquiring shells beyond b=3000 s/mm^2^, but a more comprehensive analysis would be necessary to find the optimal acquisition for DSN estimation. Structural similarity analysis could consist of an optimal similarity network via assembling several features from multiple modalities or computing several single-feature similarity networks, each sensitive to one aspect of structural similarity, for multi-network analysis [Sebenius et al., 2025].

## Conclusion

DSN analysis provides a practical and informative tool for mapping the structural organization of the human brain. It can be seamlessly integrated into most clinical dMRI neuroimaging workflows and can be accurately computed with a single shell consisting of only 30 volumes (∼3-minute acquisition). The enhanced sensitivity, biological validity, and accessibility position DSN as a promising tool for future research aimed at investigating individual brain similarity networks throughout normative and disordered processes.

## Acknowledgements

HCP-YA data was provided by the Human Connectome Project, WU-Minn Consortium (Principal Investigators: David Van Essen and Kamil Ugurbil; 1U54MH091657) funded by the 16 NIH Institutes and Centers that support the NIH Blueprint for Neuroscience Research; and by the McDonnell Center for Systems Neuroscience at Washington University. MGH-USC data was provided by the Human Connectome Project, MGH-USC Consortium (Principal Investigators: Bruce R. Rosen, Arthur W. Toga and Van Wedeen; U01MH093765) funded by the NIH Blueprint Initiative for Neuroscience Research grant; the National Institutes of Health grant P41EB015896; and the Instrumentation Grants S10RR023043, 1S10RR023401, 1S10RR019307."

## Funding Information

We acknowledge funding from DoD W81XWH-14-2-0176, NIH 5R01MH116950 and the Weill Neurohub.

## Author Information

**Radiology and Biomedical Imaging, University of California, San Francisco**

Amir Sadikov, Hannah L. Choi, Lanya T. Cai, Pratik Mukherjee

**Graduate Group in Bioengineering, University of California, San Francisco**

Amir Sadikov, Pratik Mukherjee

A.S and P.M conceived the methodology. A.S performed all analyses and drafted the manuscript. P.M supervised all analyses and contributed to writing the manuscript. H.L.C and L.T.C contributed to manuscript preparation.

## Notes

https://github.com/ucsfncl/DSN

### Competing Interest Statement

The authors have declared no competing interest.

