## Supplementary Material for "Estimating Brain Similarity Networks with Diffusion MRI"

**Figure S1: MH-USC Diffusion Similarity Networks**

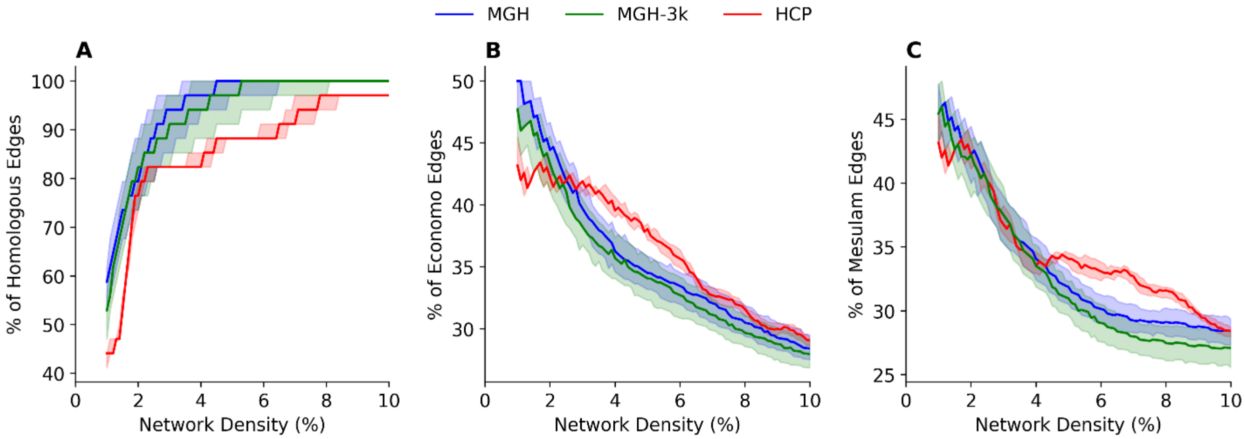


(A) the percentage of connections between regions that are homologous, (B) within the same cytoarchitectural von-Economo and Koskinas class, and (C) within the same Mesulam class over varying network densities (1% - 10%) using group-averaged DSNs derived from the HCP-YA (n=968) and MGH-USC (n=32) datasets. The shaded regions (A-C) represent the 95% confidence interval estimated by subject bootstrapping. Statistics were acquired for diffusion similarity networks computed using HCP-YA data (HCP), MGH-USC data using only b=1000 s/mm$^2$ and b=3000s/mm$^2$ shells (MGH-3k), and MGH-USC data using all shells, including b=5000 s/mm$^2$ and b=10000s/mm$^2$. DSNs were constructed using the 68 region Desikan parcellation with only cortical regions and cortico-cortical connections considered.

**Figure S2: Relating Structural Connectivity to Network Connectomes**

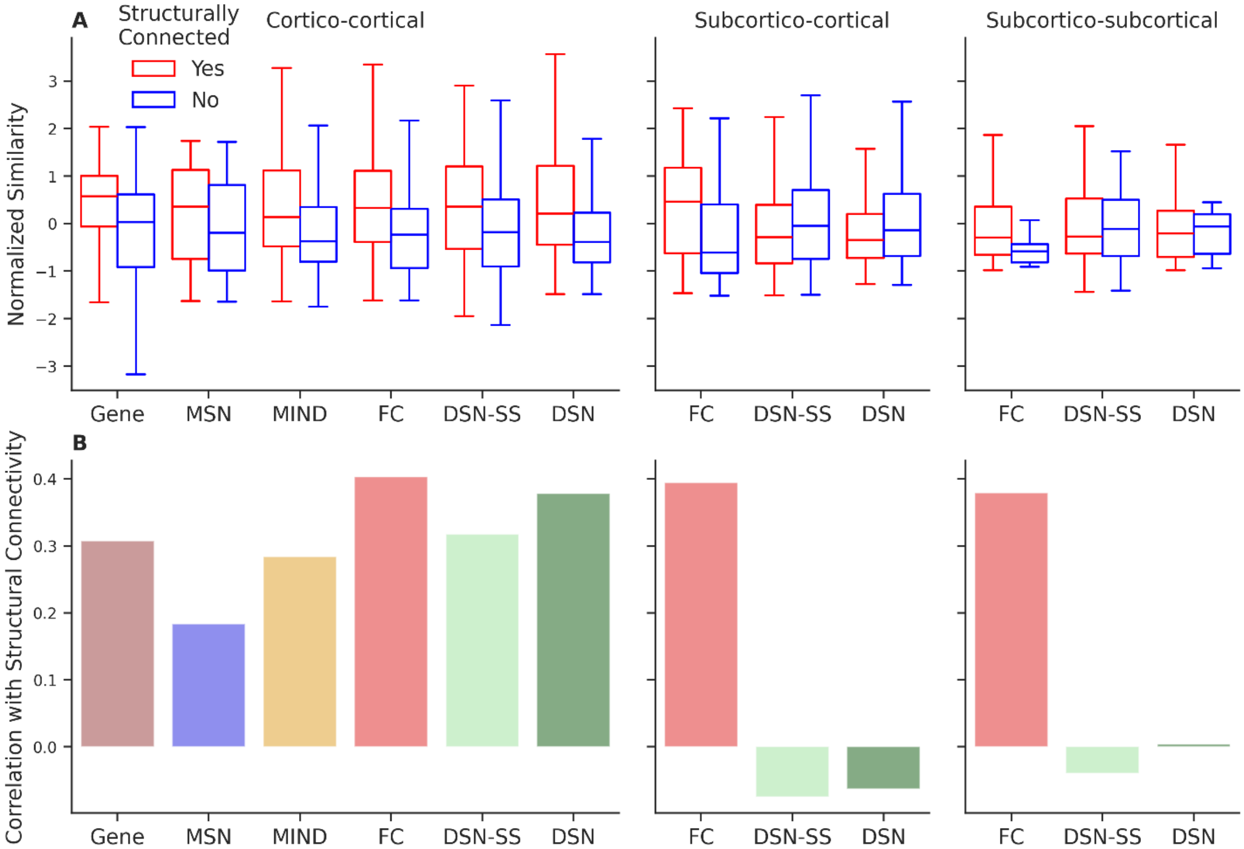


(A) The group-averaged normalized structural similarity for regions that are connected (red) vs. not connected (blue) as measured by dMRI tractography. (B) The Pearson correlation between structural similarity and other network connectomes. We compute these statistics over cortico-cortical, subcortico-cortical, and subcortico-subcortical connections for the following networks: gene expression (GENE), morphometric similarity networks (MSNs), morphometric inverse divergence (MIND) networks, functional connectivity (FC), structural connectivity (SC), and diffusion similarity networks for single-shell (DSN-SS) and multi-shell (DSN) acquisitions.

**Figure S3: Structural Similarity vs. Retrograde Tract Tracing Axonal Connectivity**

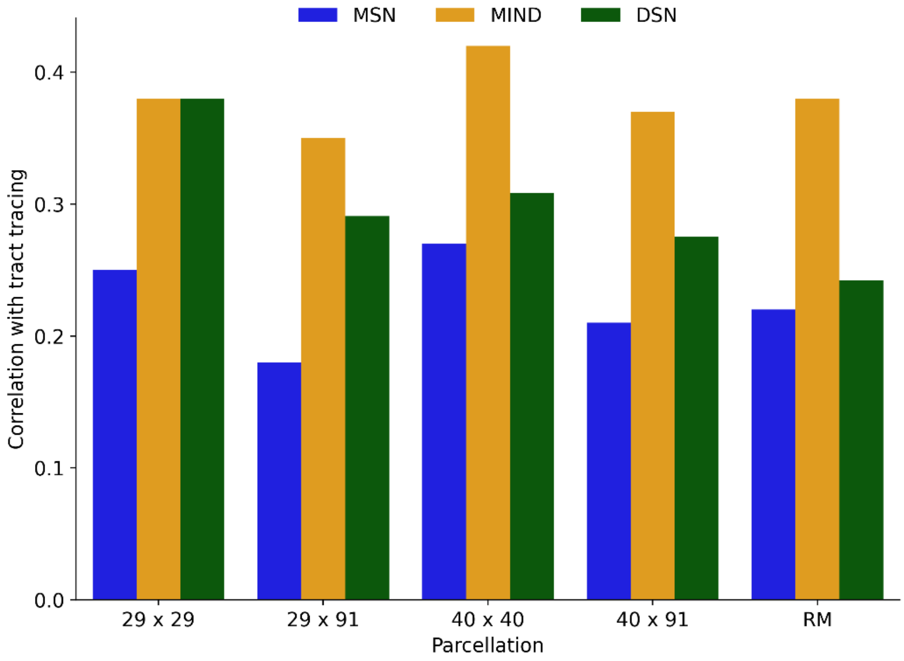


The correlation between structural similarity measured by MSN, MIND, and DSN and axonal connectivity measured via retrograde tract tracing, specifically the fraction of labeled neurons (FLNe). MIND has higher correlations with tract tracing measures than either MSN or DSN.

**Figure S4: Network Edges Sensitive to Age and Sex**

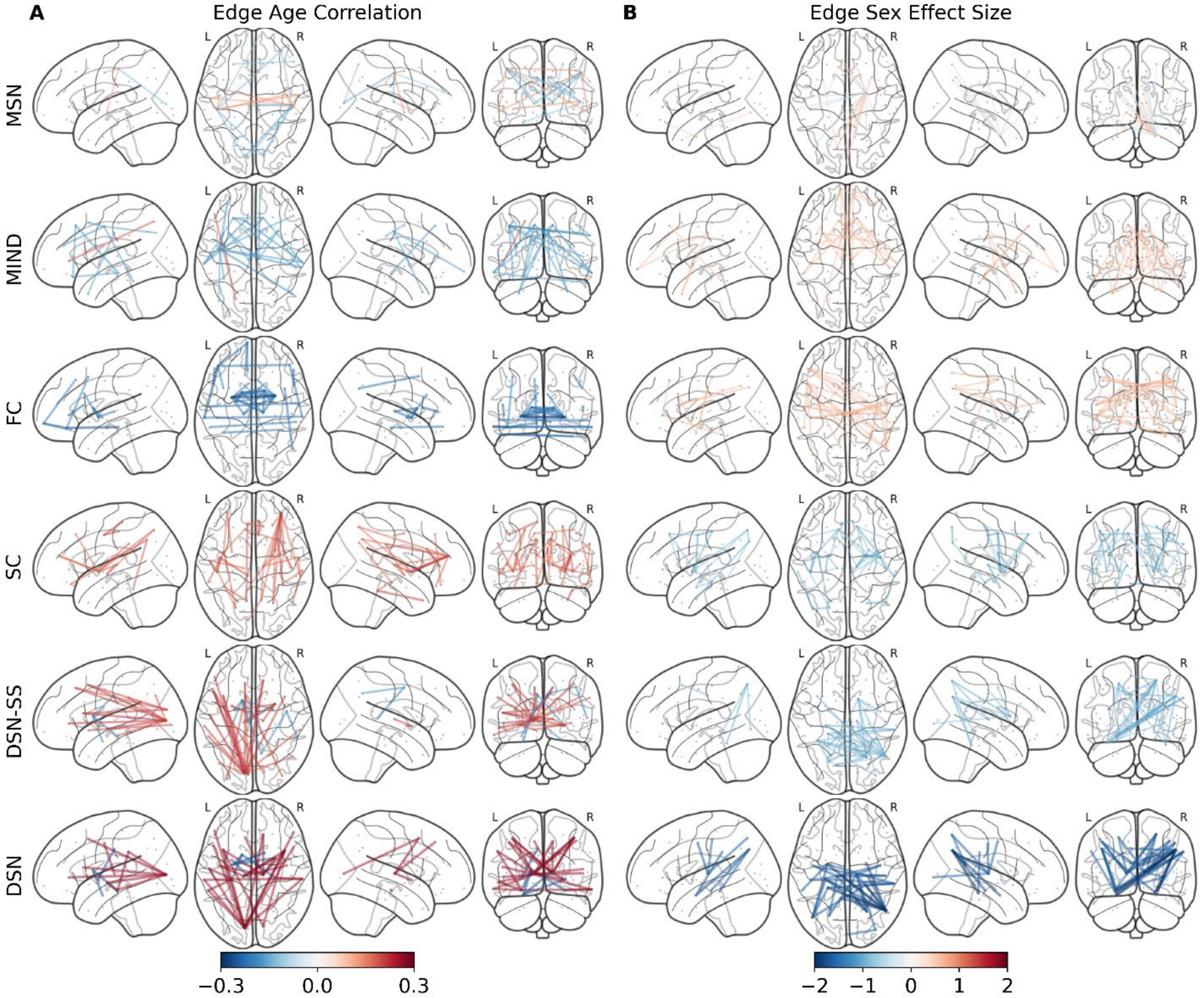


The top 1% of edges that have (A) the greatest absolute Pearson correlation coefficient with respect to age and (B) the greatest absolute effect size (male vs. female) with respect to sex for the following networks: morphometric similarity networks (MSNs), morphometric inverse divergence (MIND) networks, functional connectivity (FC), structural connectivity (SC), and diffusion similarity networks computed from single shell (DSN-SS) and multi-shell (DSN) acquisitions. Red indicates positive effect size or correlation, blue signifies negative.

**Figure S5: Network Degree Sensitive to Age and Sex**

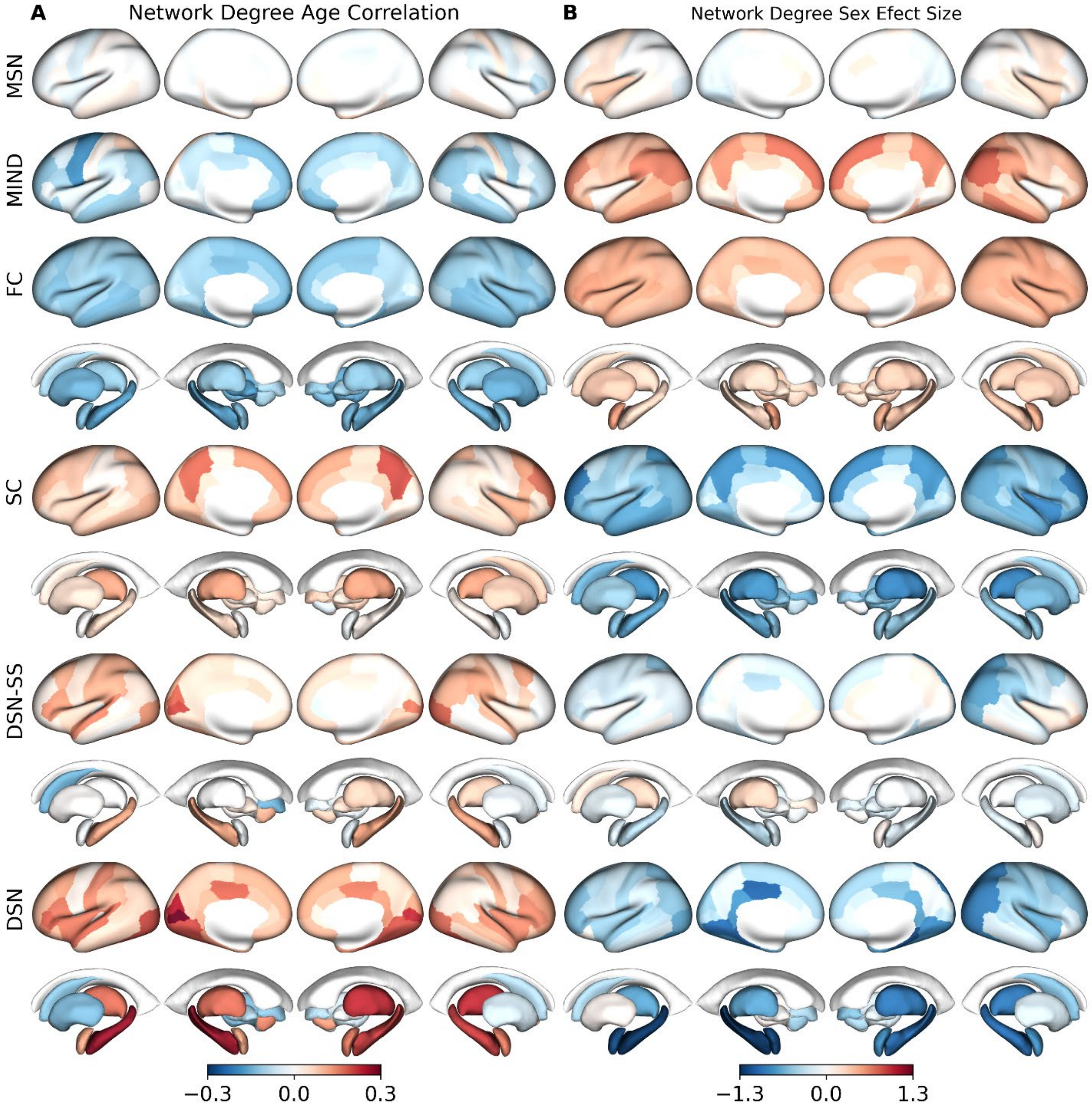


The network degree (A) Pearson correlation coefficient with respect to age and (B) effect size (male vs. female) with respect to sex for the following networks: morphometric similarity networks (MSNs), morphometric inverse divergence (MIND) networks, functional connectivity (FC), structural connectivity (SC), and diffusion similarity networks computed from single shell (DSN-SS) and multi-shell (DSN) acquisitions. Red indicates positive effect size or correlation, blue signifies negative.

**Figure S6: Correlation of Sex and Age Network Sensitivity**

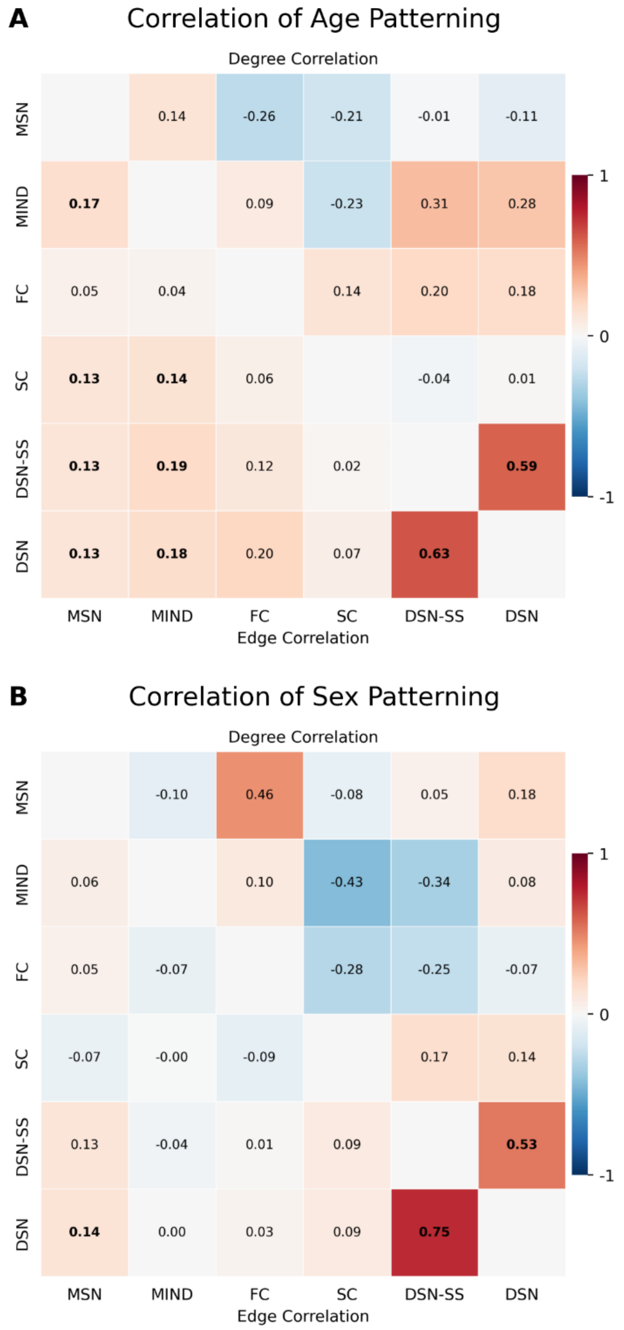


The Pearson correlation coefficient of (A) age patterning (Pearson correlation coefficients with respect to age) and (B) sex patterning (effect size: male vs. female) for edge weights (bottom/left) and network degree (top/right) between the following networks: morphometric similarity networks (MSNs), morphometric inverse divergence (MIND) networks, functional connectivity (FC), structural connectivity (SC), and diffusion similarity networks computed from single shell (DSN-SS) and multi-shell (DSN) acquisitions. Red indicates positive effect size or correlation, blue signifies negative. Bolded values indicate statistical significance following a two-sided spin test with false discovery rate correction across all correlations.

**Figure S7: Mapping Subcortical Network Connectome Similarity**

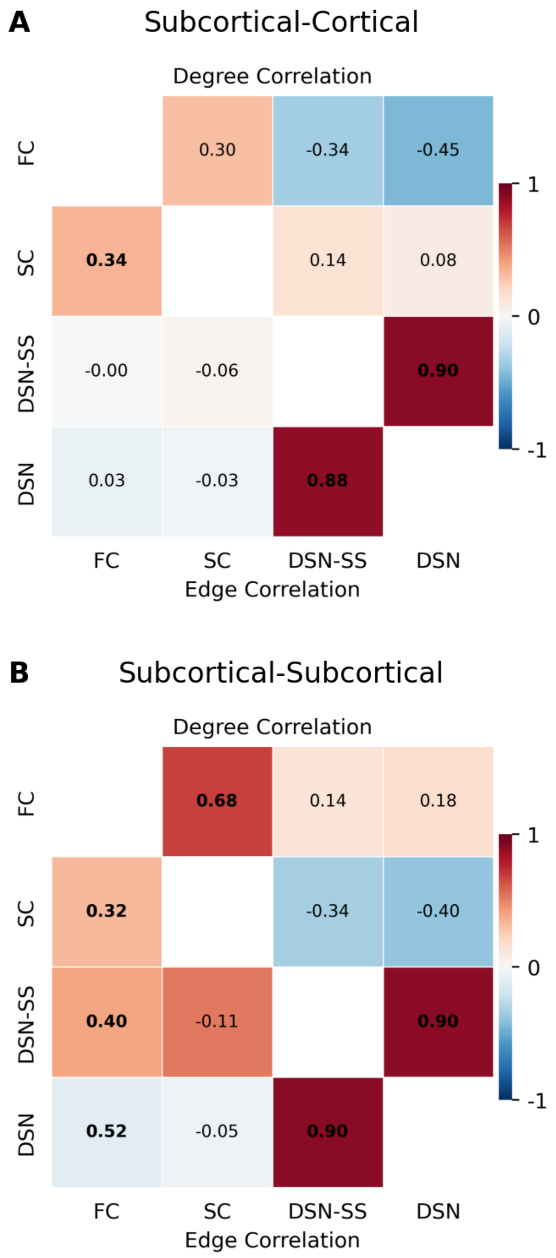


The Pearson correlation coefficient between group-averaged network edge weights (bottom/left) and node centrality (top/right) for the (A) subcortical-cortical and (B) subcortical-subcortical portions of the connectome for the following networks: functional connectivity (FC), structural connectivity (SC), and diffusion similarity networks computed from single shell (DSN-SS) and multi-shell (DSN) acquisitions. Red indicates positive correlation, blue signifies negative. Bolded values indicate statistical significance following a two-sided spin test with false discovery rate correction across all correlations.

**Figure S8: Comparing Network Connectome Laplacian Eigenspaces**

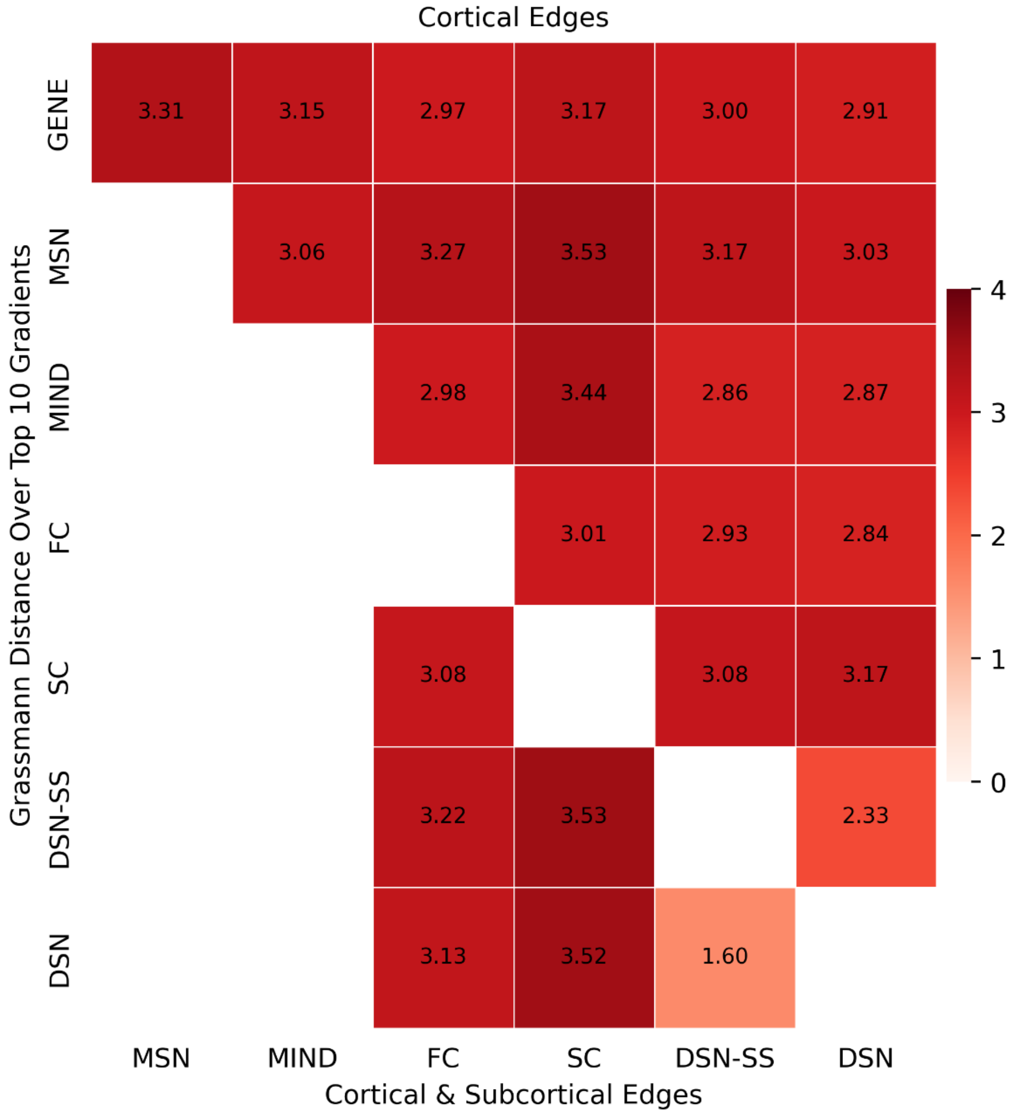


The Grassmann distance between group-averaged network connectome Laplacian eigenspaces over only cortical edges (top/right) and over cortical and subcortical edges (bottom/left) for the following networks: gene expression (GENE), morphometric similarity networks (MSN), morphometric inverse divergence (MIND), functional connectivity (FC), structural connectivity (SC), and diffusion similarity networks computed from single shell (DSN-SS) and multi-shell (DSN) acquisitions. Laplacian eigenspaces were derived by selecting the top ten gradients from the normalized Laplacian embedding of each group-averaged network connectome. The stationary gradient was excluded.

**Figure S9: Edge Heritability across Network Connectomes**

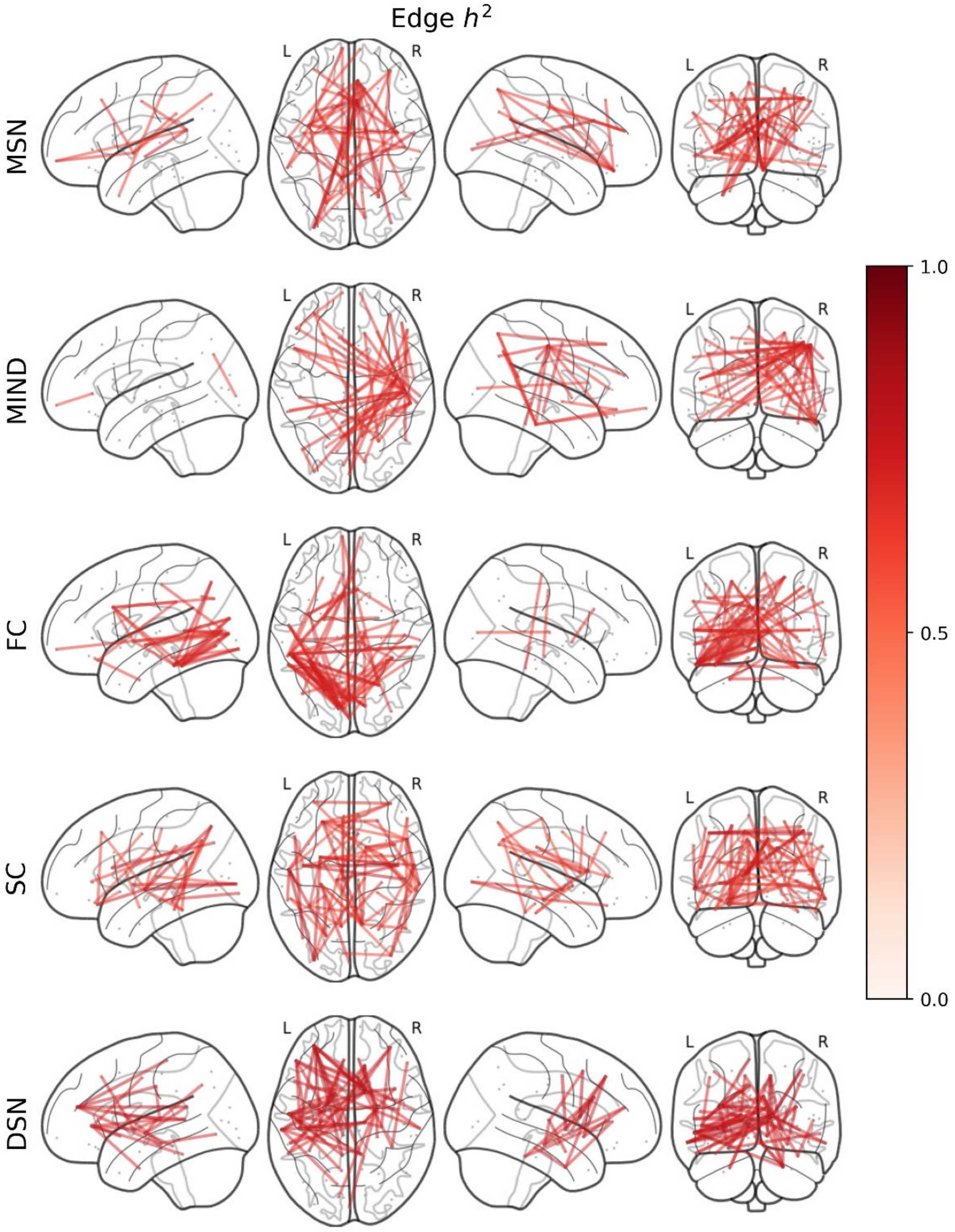


The top 1% of edge weight heritability ($h^2$) across the following networks: morphometric similarity networks (MSNs), morphometric inverse divergence (MIND) networks, functional connectivity (FC), structural connectivity (SC), and diffusion similarity networks (DSNs).

**Figure S10: Degree Heritability across Network Connectomes**

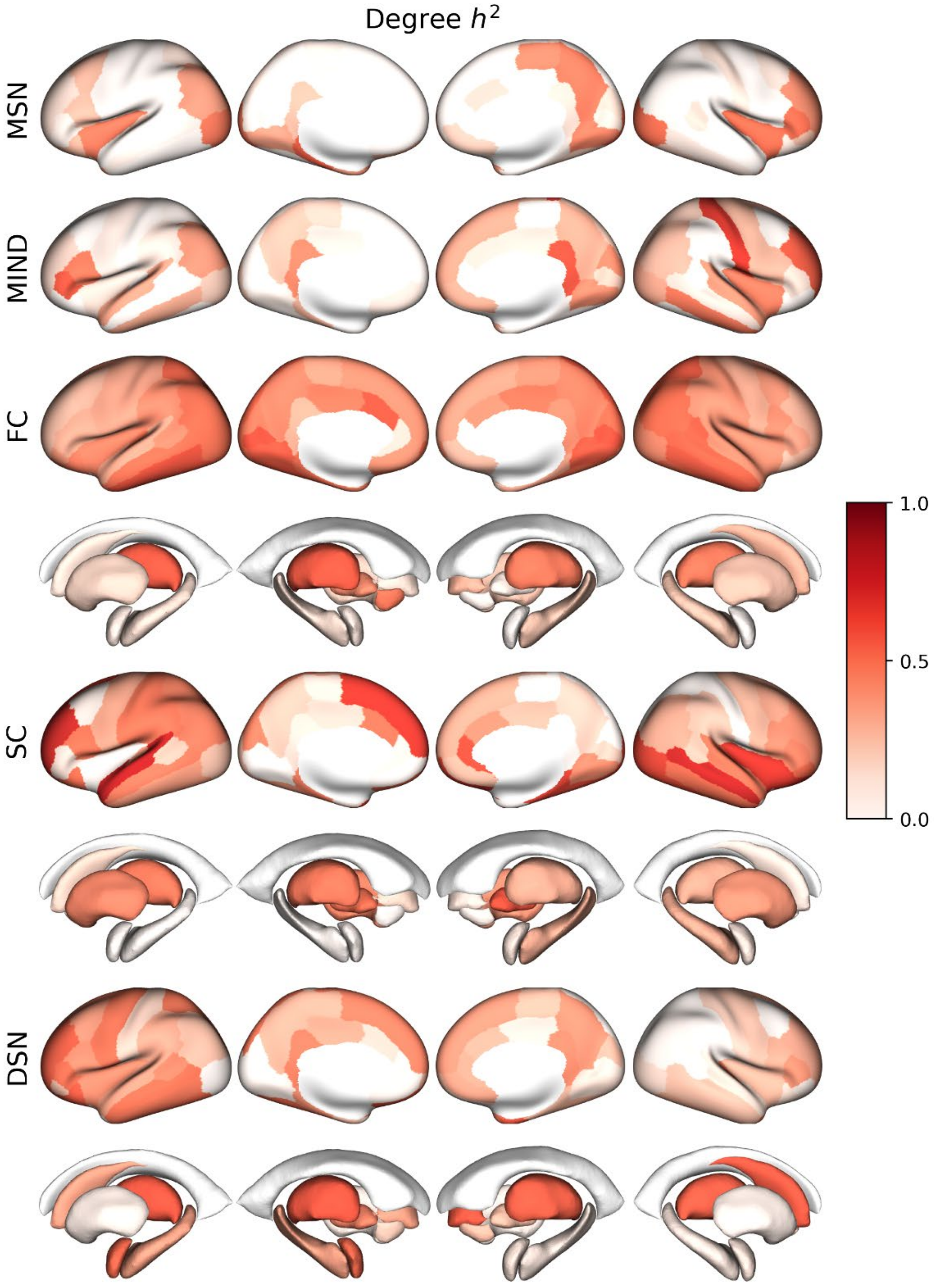


The network degree heritability ($h^2$) across the following networks: morphometric similarity networks (MSNs), morphometric inverse divergence (MIND) networks, functional connectivity (FC), structural connectivity (SC), and diffusion similarity networks (DSNs).

**Figure S11: Heritability of Structurally Connected Regions**

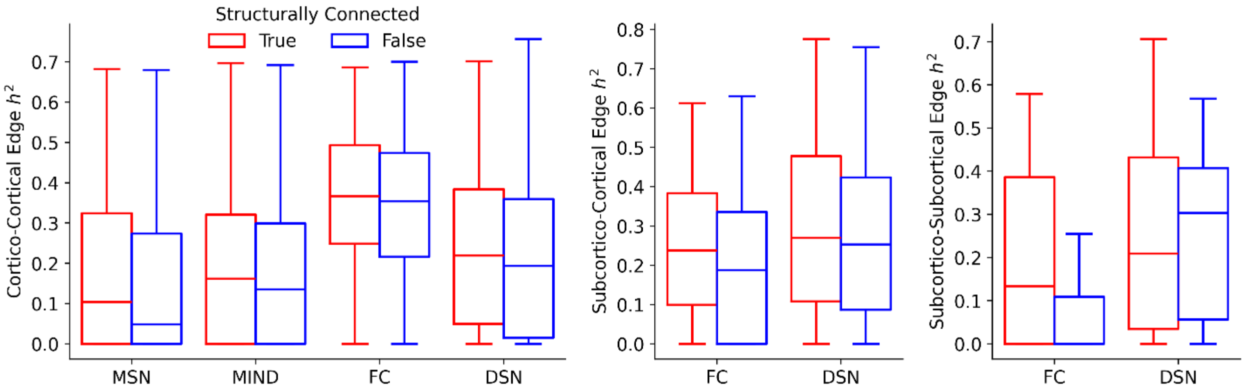


The edge weight heritability ($h^2$) across cortico-cortical, subcortico-cortical, and subcortico-subcortical connections. Structural similarity, as measured by DSN, between structurally connected regions is more heritable than between those that are not structurally connected for cortico-cortical (t=1.9, p=0.058) and subcortico-cortical connections (t=2.44, p=0.015). Statistical significance was assessed using two-sided independent t-tests with unequal variances. These results were similar for the other networks.

### Figure S12: Mapping Network Degree Heritability across the Cortex

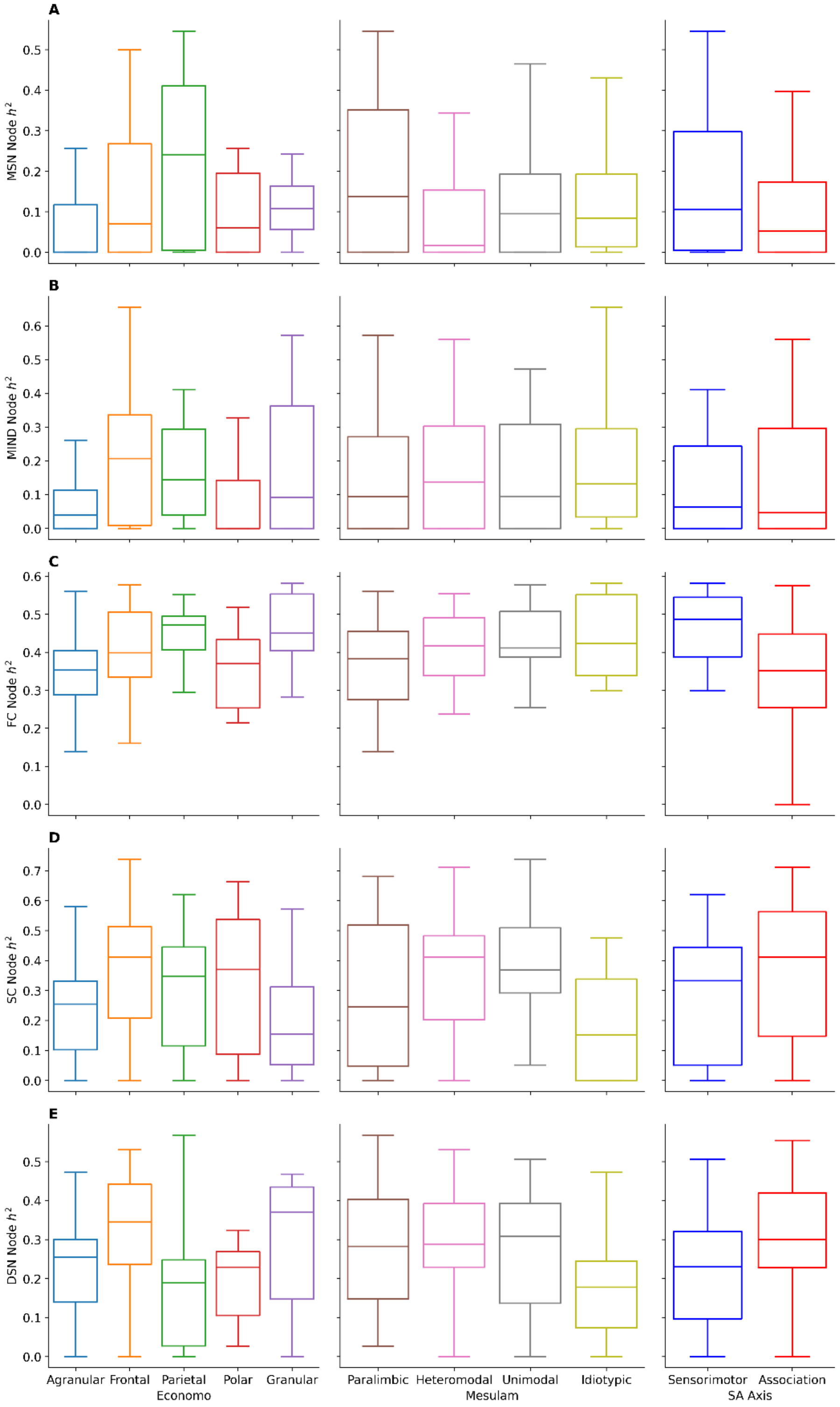


The network degree heritability across the von Economo and Koskinas (left), Mesulam (center), and sensorimotor-association (right) parcellations for (A) morphometric similarity networks (MSNs), (B) morphometric inverse divergence (MIND), (C) functional connectivity (FC), (D) structural connectivity (SC), and (E) diffusion similarity networks (DSNs).

**Figure S13: Correlation of Heritability Patterning**

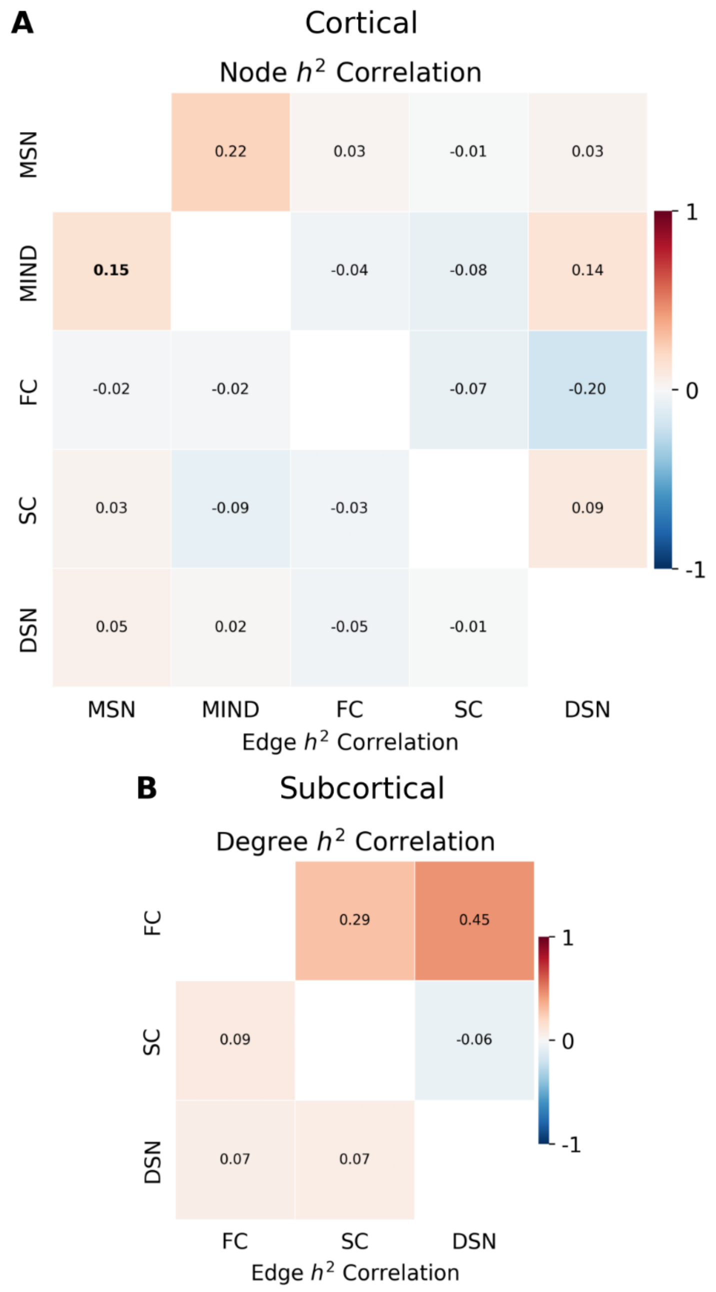


The Pearson correlation coefficient of heritability patterning for (A) cortical and (B) subcortical nodes for edge weights (bottom/left) and network degree (top/right) between the following networks: morphometric similarity networks (MSNs), morphometric inverse divergence (MIND) networks, functional connectivity (FC), structural connectivity (SC), and diffusion similarity networks (DSNs). Red indicates positive effect size or correlation, blue signifies negative. Bolded values indicate statistical significance following a two-sided spin test with false discovery rate correction across all correlations.

**Figure S14: Edge Weight vs. Heritability**

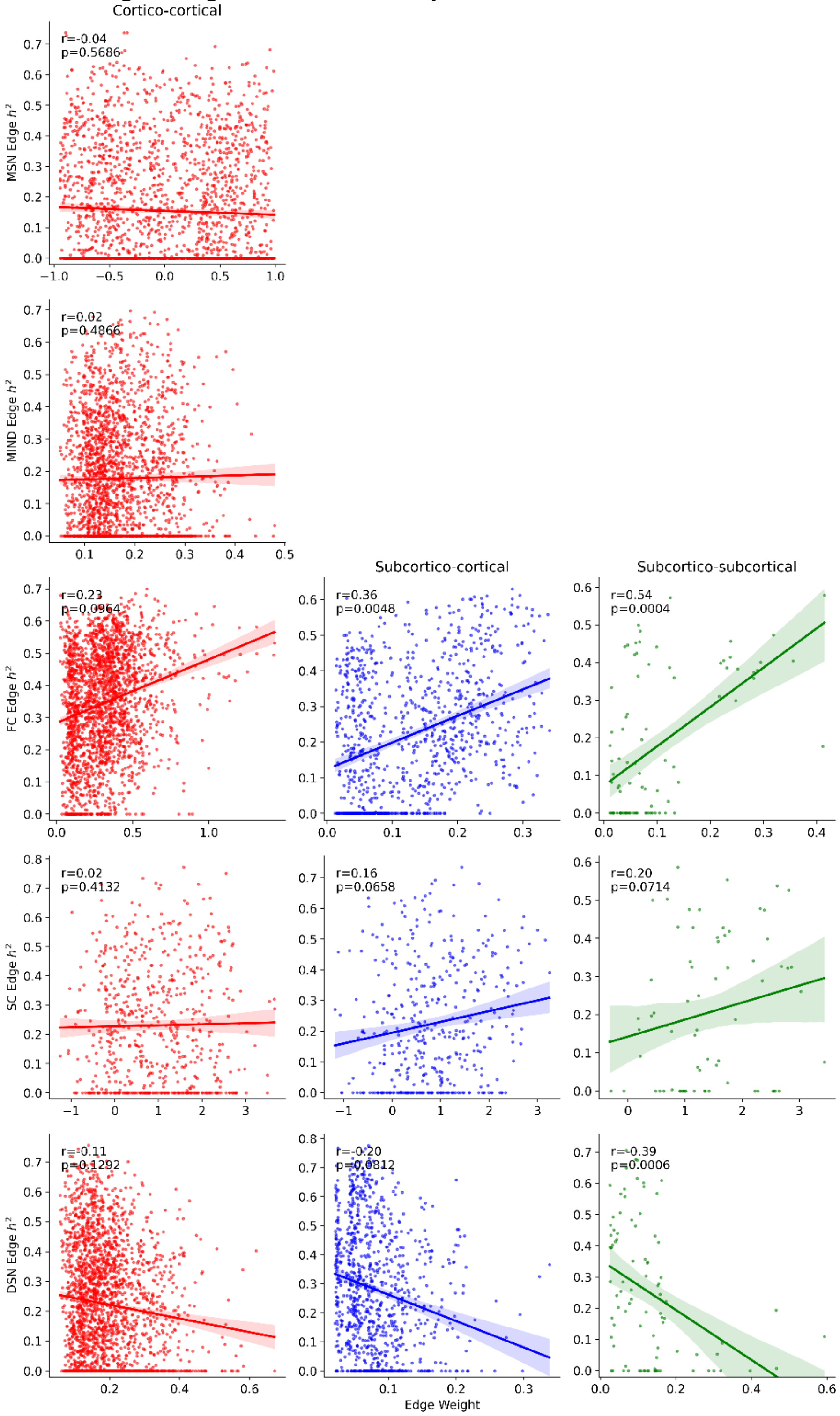


The correlation between edge weight and heritability ($h^2$) across cortico-cortical (red), subcortico-cortical (blue), and subcortico-subcortical (green) connections for the following networks: morphometric similarity networks (MSNs), morphometric inverse divergence (MIND) networks, functional connectivity (FC), structural connectivity (SC), and diffusion similarity networks computed from single shell (DSN-SS) and multi-shell (DSN) acquisitions.

**Figure S15: Network Degree vs. Heritability**

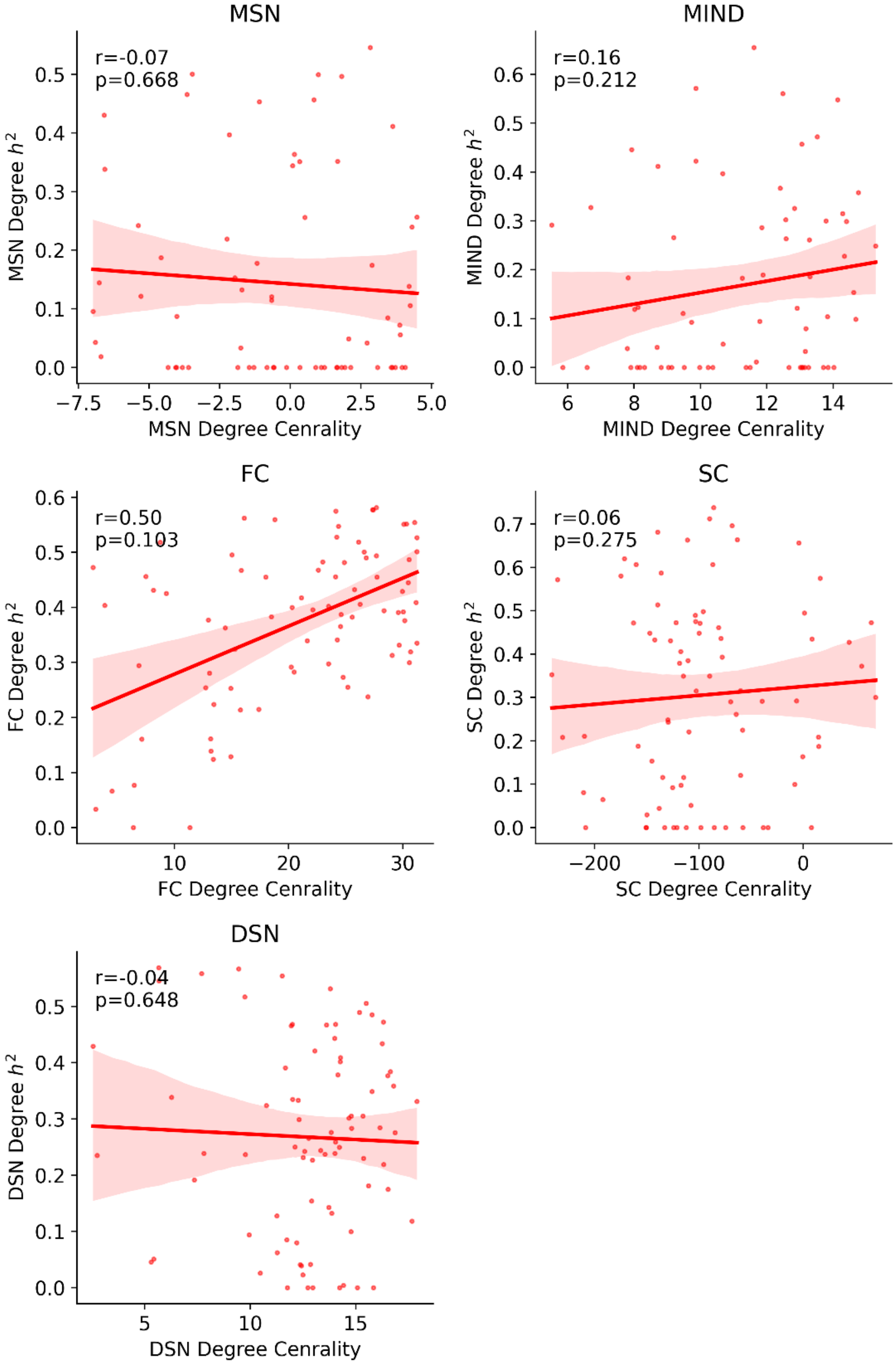


The correlation between network degree and heritability ($h^2$) for the following networks: morphometric similarity networks (MSNs), morphometric inverse divergence (MIND) networks, functional connectivity (FC), structural connectivity (SC), and diffusion similarity networks (DSNs).

**Table S1: Predicting Age, Cognition, and Sex with Edge Weight**

| **Coefficient of Determination & AUC-ROC** | | | | | |
|---|---|---|---|---|---|
| Network | Age ($R^2$) | Fluid Cognition ($R^2$) | Crystalized Cognition ($R^2$) | Total Cognition ($R^2$) | Sex (AUC-ROC) |
| MSN | 0.001 +/- 0.026 | 0.009 +/- 0.025 | 0.029 +/- 0.022 | 0.022 +/- 0.028 | 0.794 +/- 0.031 |
| MIND | 0.065 +/- 0.036 | 0.019 +/- 0.025 | 0.084 +/- 0.028 | 0.074 +/- 0.033 | 0.911 +/- 0.017 |
| FC | 0.186 +/- 0.049 | **0.055 +/- 0.047** | 0.119 +/- 0.040 | 0.115 +/- 0.048 | 0.930 +/- 0.015 |
| SC | 0.210 +/- 0.043 | 0.054 +/- 0.026 | 0.101 +/- 0.044 | 0.080 +/- 0.039 | 0.971 +/- 0.009 |
| DSN-SS | 0.263 +/- 0.054 | 0.023 +/- 0.029 | 0.118 +/- 0.041 | 0.088 +/- 0.042 | 0.964 +/- 0.012 |
| DSN | **0.325 +/- 0.049** | 0.049 +/- 0.034 | **0.148 +/- 0.043** | **0.123 +/- 0.037** | **0.983 +/- 0.006** |
| **One-sided paired t-test p-values with respect to DSN** | | | | | |
| Network | Age (p) | Fluid Cognition (p) | Crystalized Cognition (p) | Total Cognition (p) | Sex (p) |
| MSN | **9.5e-78** | **3.6e-18** | **3.2e-49** | **2.5e-46** | **6.7e-82** |
| MIND | **4.0e-73** | **1.4e-15** | **2.5e-29** | **2.4e-23** | **1.3e-66** |
| FC | **1.2e-39** | 1.0 | **2.8e-09** | 0.081 | **3.4e-56** |
| SC | **4.2e-34** | 1.0 | **3.6e-17** | **1.0e-20** | **1.1e-21** |
| **One-sided paired t-test p-values with respect to DSN-SS** | | | | | |
| Network | Age (p) | Fluid Cognition (p) | Crystalized Cognition (p) | Total Cognition (p) | Sex (p) |
| MSN | **3.2e-66** | **7.5e-05** | **2.8e-38** | **2.7e-26** | **4.2e-76** |
| MIND | **1.4e-55** | 0.12 | **4.2e-12** | **7.1e-04** | **3.8e-52** |
| FC | **1.2e-19** | 1.0 | 0.61 | 1.0 | **1.1e-33** |
| SC | **1.3e-13** | 1.0 | **3.3e-04** | **0.029** | 1.0 |

The statistics for this table are computed for the network edge weights. (Top) The average and standard deviation of the coefficients of determination ($R^2$) for the prediction of age, fluid cognition, and total cognition and area under the receiver operator curve (AUC-ROC) for the prediction of sex across one-hundred cross-validation folds (five-fold repeated cross validation; 20 repeats). The p-values from one-sided paired t-tests following false discovery rate correction with respect to (Middle) the multi-shell diffusion similarity network (DSN) and (Bottom) sub-sampled single-shell diffusion similarity network (DSN-SS). DSN as well as DSN-SS are significantly more predictive than other network connectomes over the edge weights in most cases.

**Table S2: Predicting Age, Cognition, and Sex with Degree Centrality**

| **Coefficient of Determination & AUC-ROC** | | | | | |
|---|---|---|---|---|---|
| Network | Age ($R^2$) | Fluid Cognition ($R^2$) | Crystalized Cognition ($R^2$) | Total Cognition ($R^2$) | Sex (AUC-ROC) |
| MSN | -0.012 +/- 0.014 | 0.011 +/- 0.020 | 0.008 +/- 0.019 | 0.013 +/- 0.021 | 0.731 +/- 0.034 |
| MIND | 0.042 +/- 0.026 | 0.008 +/- 0.021 | 0.053 +/- 0.024 | 0.046 +/- 0.028 | 0.778 +/- 0.028 |
| FC | 0.050 +/- 0.030 | -0.004 +/- 0.022 | 0.017 +/- 0.027 | 0.008 +/- 0.022 | 0.782 +/- 0.027 |
| SC | 0.023 +/- 0.025 | 0.006 +/- 0.022 | 0.019 +/- 0.023 | 0.021 +/- 0.026 | 0.857 +/- 0.024 |
| DSN-SS | 0.116 +/- 0.043 | 0.005 +/- 0.023 | 0.089 +/- 0.037 | 0.063 +/- 0.030 | 0.881 +/- 0.021 |
| DSN | **0.185 +/- 0.048** | **0.025 +/- 0.030** | **0.115 +/- 0.043** | **0.089 +/- 0.037** | **0.952 +/- 0.012** |
| **One-sided paired t-test p-values with respect to DSN** | | | | | |
| Network | Age (p) | Fluid Cognition (p) | Crystalized Cognition (p) | Total Cognition (p) | Sex (p) |
| MSN | **1.1e-62** | **4.3e-06** | **4.2e-41** | **1.2e-38** | **2.3e-85** |
| MIND | **4.6e-51** | **2.3e-08** | **5.1e-25** | **5.1e-21** | **4.2e-81** |
| FC | **7.3e-44** | **5.8e-16** | **1.0e-42** | **1.8e-38** | **3.0e-81** |
| SC | **2.7e-56** | **6.9e-10** | **2.1e-40** | **7.1e-39** | **7.7e-65** |
| **One-sided paired t-test p-values with respect to DSN-SS** | | | | | |
| Network | Age (p) | Fluid Cognition (p) | Crystalized Cognition (p) | Total Cognition (p) | Sex (p) |
| MSN | **1.0e-50** | 1.0 | **3.5e-35** | **5.8e-30** | **6.7e-62** |
| MIND | **1.7e-29** | 1.0 | **4.0e-15** | **3.8e-08** | **1.7e-56** |
| FC | **2.7e-23** | **2.2e-04** | **8.7e-34** | **6.3e-29** | **1.8e-53** |
| SC | **9.3e-41** | 1.0 | **1.5e-33** | **3.7e-27** | **1.2e-13** |

The statistics for this table are computed for the network degree centralities. (Top) The average and standard deviation of the coefficients of determination ($R^2$) for the prediction of age, fluid cognition, and total cognition and area under the receiver operator curve (AUC-ROC) for the prediction of sex across one-hundred cross-validation folds (five-fold repeated cross validation; 20 repeats). The p-values from one-sided paired t-tests following false discovery rate correction with respect to (Middle) the multi-shell diffusion similarity network (DSN) and (Bottom) sub-sampled single-shell diffusion similarity network (DSN-SS). DSN as well as DSN-SS are significantly more predictive than most other network connectomes over the node centralities weights in most cases.
